# Reconstitution of Ras-PI3Kγ membrane communication and feedback using light-induced signaling inputs

**DOI:** 10.64898/2026.02.17.706477

**Authors:** Sophia Doerr, Andres Olavarrieta-Colasurdo, Scott D. Hansen

## Abstract

Biochemical reactions involving phosphatidylinositol phosphate (PIP) lipids and small GTPases serve essential roles in signal transduction immediately downstream of cell surface receptor activation. Interconnected positive and negative feedback loops link these distinct classes of reactions and allow for the rapid establishment of the steep and precisely oriented intracellular activity gradients that are hallmarks of cell polarity. Although genetic approaches have identified numerous molecules that potentially regulate feedback between GTPases and PIP lipids in cells, it is unclear how different types of feedback modulate the spatiotemporal dynamics of membrane signaling reactions. Here, we reconstitute communication and feedback between Ras GTPase and phosphatidylinositol 3-kinase gamma (PI3Kγ)-mediated generation of PIP3 on supported membranes. We employ light-induced membrane recruitment of a guanine nucleotide exchange factor (GEF) to rapidly and reversibly shift steady-state conditions and monitor corresponding Ras activation and PIP3 production in time and space. Alone, the Ras-PI3Kγ module exhibits weak and transient activation due to global inhibitors. However, the introduction of GEF-mediated positive feedback drives local amplification of Ras activation and PIP3 generation, resulting in a wave of activity that propagates across the membrane surface. Spatial coupling between activated Ras and PIP3 molecules depends on their respective diffusion coefficients and the species involved in the feedback circuit. This work identifies activation thresholds, membrane diffusion, and feedback architecture as key factors that determine how small GTPases and PIP lipid modifying enzymes amplify membrane signaling reactions in the presence of global inhibitors.

## INTRODUCTION

To execute their specialized functions, cells must sense and rapidly respond to environmental signals. Two classes of molecules that are ubiquitously important for signal transduction at the plasma membrane of eukaryotic cells are phosphatidylinositol phosphate (PIP) lipids and small GTPases (**Fig. 1a, b**) ^1,2^. Together, these molecules function downstream of numerous plasma membrane receptors to control processes such as chemotaxis, neurogenesis, and asymmetric cell division. The abundance and spatial distribution of PIP lipids and activated small GTPases are controlled by an interconnected network of positive and negative feedback loops ^3–6^. Positive feedback in plasma membrane-associated signaling events functions to amplify external signals and break spatial symmetry through the rapid generation of molecular concentration gradients inside cells ^7–9^. Currently, it is challenging to decipher how specific feedback circuits control the spatiotemporal dynamics of individual signaling modules in vivo due to redundancy and crosstalk within signaling networks. To understand feedback circuitry in signal processing, a bottom-up reductionist biochemistry approach is necessary to complement the large body of existing cell biological research.

**Figure 1.**
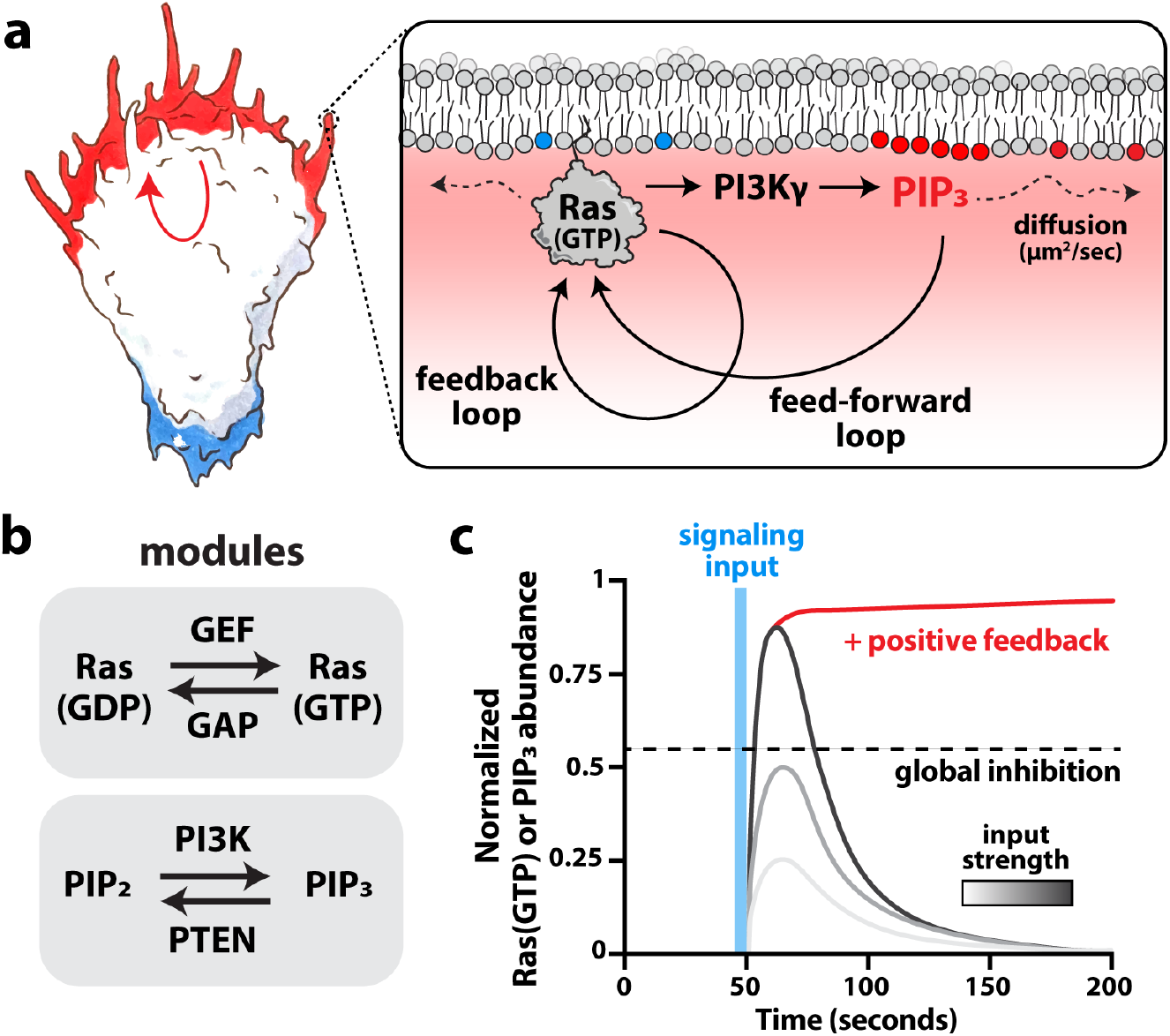
Positive feedback allows for the amplification of Ras and PIP_3_ signaling. **(a)** Membrane-associated GTPase activation and PIP lipid phosphorylation regulates early signaling reactions that govern cell polarity. Active Ras GTPase promotes PI3Kγ membrane localization which results in PIP_3_ generation. GEF-mediated positive feedback and feed-forward loops are thought to amplify Ras and PIP_3_ signaling in vivo. **(b)** GTPases in the Ras superfamily are activated by guanine nucleotide exchange factors (GEFs) and inactivated by GTPase activating proteins (GAPs). Phosphatidylinositol phosphate (PIP) lipids are interconverted by lipid kinases and phosphatases (e.g. PI3K and PTEN). **(c)** In vivo, membrane signaling reactions are often transient and reversible due to global inhibition and negative feedback. Positive feedback drives signal amplification and a sustained cellular response.

Phosphatidylinositol (3,4,5)-trisphosphate (PIP_3_) lipids are essential signaling molecules that are generated primarily through the phosphorylation of PIP_2_ by class I phosphatidylinositol 3-kinases at the plasma membrane ^10^. Across cell types, the production of PIP_3_ plays an instructive role in building cortical dendritic actin networks ^11,12^. In hematopoietic cells, the phosphatidylinositol 3-kinase gamma (PI3Kγ) isoform serves critical functions in regulating chemotaxis, phagocytosis, cytokine production, and the generation of reactive oxygen species (ROS) ^13,14^. PI3Kγ is activated at the plasma membrane downstream of G-protein coupled receptors (GPCRs) and the GTP-bound form of Ras GTPase ^15–17^. Since the Ras-PI3Kγ signaling module is broadly important for immune cell function, dysregulation of this pathway has been linked to numerous inflammatory diseases and cancers ^13,18,19^. Studying the mechanisms that control communication between PI3Kγ and Ras GTPase has far-reaching implications for our understanding of cell signaling.

Biochemical reconstitutions of PIP lipid modifying enzymes and small GTPases have historically relied on solution-based measurements that fail to capture the spatial organization and membrane association dynamics of these proteins. However, recent advances in fluorescence microscopy and supported lipid bilayer (SLB) technology have enabled researchers to directly visualize the membrane binding and activity of many peripheral membrane proteins with single molecule resolution ^20–22^. Here, we reconstitute Ras and PI3Kγ activation reactions on SLBs and observe their spatial evolution using Total Internal Reflection Fluorescence (TIRF) Microscopy. Despite these technical innovations, kinetic measurements still often rely solely on simply monitoring the conversion of substrate to product in a single, irreversible direction. By contrast, cell signaling reactions involving PIP lipid modifying enzymes and small GTPases are typically transient and reversible due to global inhibitors and product-induced negative feedback loops (**Fig. 1c**)^23^. It is the dynamic interconversion between substrate and product that gives rise to unique emergent properties in cells, such as bistability ^24^, cortical excitability ^8^, and polarity ^25^.

To mimic the spatiotemporal dynamics of cell signaling events we utilize the improved Light-Inducible Dimer (iLID) system ^26^ to reversibly control the localization of peripheral membrane binding proteins that shape the membrane signaling landscape. This approach allows us to rapidly shift steady-state reaction conditions and visualize the dynamic response of the Ras-PI3Kγ signaling module. By combining this light-inducible system with spatial control granted by a digital micromirror device (DMD), we locally trigger biochemical reactions and observe how secondary messengers respond in time and space. Introducing positive feedback loops mediated by guanine nucleotide exchange factors (GEFs) allows us to determine conditions under which the Ras-PI3Kγ signaling module can overcome global inhibition imposed by a GTPase activating protein (GAP) and the lipid phosphatase PTEN (**Fig. 1a, b**). Positive feedback results in the spatiotemporal amplification of both Ras activation and PI3Kγ-dependent production of PIP_3_. This work establishes a novel experimental framework for systematically studying how feedback enables membrane signaling reactions to overcome global inhibition and drive spatial patterning programs. Deciphering how the interconversion of PIP lipids and small GTPases function in concert provides insight into the mechanisms that govern signal transduction across cell types.

## RESULTS

### Spatial control of light-induced membrane recruitment

To study communication and feedback between small GTPases and PIP lipid modifying enzymes, we adapted the improved Light-Inducible Dimer (iLID) system ^26^ to spatially and temporally control the localization of purified proteins on supported lipid bilayers (SLBs). For these experiments, purified his_10_-mNeonGreen-iLID (referred to as iLID) was stably tethered to SLBs using nickel-chelating lipids and visualized by TIRF microscopy (**Fig. 2a**). In the absence of 488 nm (blue) light, iLID adopts a conformation that sterically occludes its SsrA peptide from interacting with its binding partner, SspB (stringent response protein B) (**Fig. 2a**) ^26^. Although iLID is considered “closed” in the dark, it is still conformationally dynamic and displays a weak affinity for some SspB variants ^26^. To measure the degree of light-independent association between the two proteins, we compared the membrane absorption kinetics and relative equilibrium membrane densities of three previously isolated SspB variants (i.e. nano, micro, and milli) ^27^ in the presence of membrane-tethered iLID (**Fig. 2b**). In the absence of 488 nm light, equilibrium membrane density of the high-affinity SspB(nano) variant was ≥10-fold greater than those of SspB(micro) and SspB(milli) (**Fig. 2b**).

**Figure 2.**
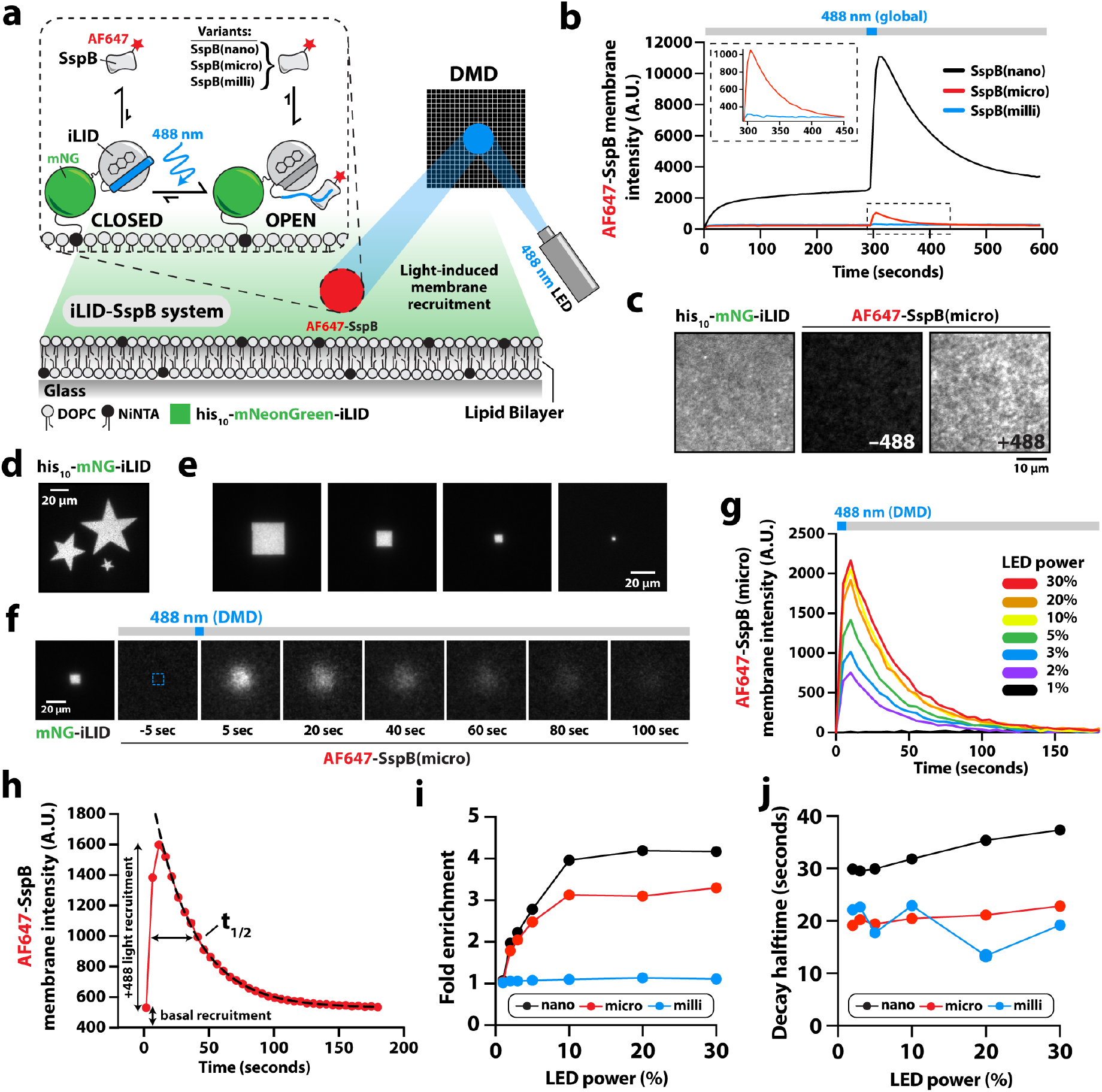
Spatially controlled light-induced heterodimerization of iLID and SspB on reconstituted membranes. **(a)** Schematic cartoon showing his_10_-mNG-iLID tethered to a supported lipid bilayer (SLB) via a NiNTA lipid. In the dark, its binding partner, AF647-SspB, is predominantly in solution. A digital micromirror device (DMD) can be used to spatially control the light-induced activation of membrane-tethered iLID and AF647-SspB membrane recruitment. **(b)** SspB variants (nano, micro, and milli) have distinct membrane absorption properties in the absence of 488 nm light and responses to activating light. All SspB variants exponentially dissociate from the membrane in the dark. **(c)** Representative TIRF-M images showing SLB localization of iLID and AF647-SspB(micro) in the absence and presence of uniform 488 nm light. **(d)** TIRF-M image showing DMD-generated patterns of 488 nm-excited iLID on SLBs. **(e)** TIRF-M images showing DMD-generated shapes (100, 25, 4, and 1 µm^2^) of 488 nm-excited iLID. **(f)** Montage of TIRF-M images showing the time-dependent change in AF647-SspB(micro) membrane localization following DMD-guided activation of iLID. **(g)** Kinetic traces showing AF647-SspB(micro) membrane recruitment and dissociation measured over a range of LED powers (0-50 μW, see **Supplemental Fig. 2a**). **(h)** Representative kinetic trace of AF647-SspB(micro) membrane recruitment and dissociation. Fold membrane enrichment was calculated using (basal recruitment + 488 nm light recruitment)/basal recruitment. Fitting to non-linear one phase decay curves (dashed line) were used to calculated signal decay halftime. **(i)** Quantification of AF647-SspB(nano, micro, and milli) fold membrane enrichment as a function of LED power. **(j)** Quantification of AF647-SspB(nano, micro, and milli) signal decay halftime as a function of LED power. **(b-j)** Membrane composition: 94% DOPC, 2% NiNTA (+iLID conjugated), 2% MCC-PE, and 2% PIP_2_ lipids.

After allowing the SspB variants to reach equilibrium in the presence of membrane-tethered iLID, we compared their light-induced membrane recruitment. For these experiments, we used a single, 50 ms pulse of 488 nm light (∼50 µW, **Supplementary Fig. 2a**) to shift iLID into the “open” conformation and allow it to associate with SspB. In all cases, this resulted in an immediate increase in AF647-SspB membrane localization that peaked within 10 seconds of the 488 nm activating signal (**Fig. 2b, c**). SspB(nano) displayed the highest total membrane recruitment, SspB(micro) exhibited a modest increase in membrane intensity, and the change in the SspB(milli) signal was nearly undetectable. All three SspB variants dissociated from bilayers with exponential kinetics based on the reversion of iLID to the “closed”, dark-state conformation (**Fig. 2b**).

To mimic the polarized membrane signaling that occurs in vivo, we established an approach to spatially control the blue light-induced activation of membrane-tethered iLID on SLBs using a digital micromirror device (DMD) (**Fig. 2a, d**). For different 488-nm illumination patterns, we visualized the corresponding SspB(micro) membrane localization and defined the limits of our ability to detect iLID-SspB heterodimerization (**Fig. 2e** and **Supplementary Fig. 3a, b**). For all light patterns, lateral membrane diffusion of iLID caused SspB(micro) membrane localization to expand slightly beyond the initially defined region of activation (**Fig. 2f**). We could repeatedly activate iLID in the same membrane region without observing a decrease in SspB recruitment (**Supplementary Fig. 4a**). Using spatially patterned light, we next sought to tune SspB membrane recruitment by modulating the intensity of blue light used to activate iLID. Visualization of SspB membrane recruitment as a function of blue light intensity revealed that iLID was maximally activated in the presence of light at 10% LED power (∼15 µW, **Fig. 2g**). Lower levels of activating blue light resulted in less SspB membrane recruitment (**Fig. 2g** and **Supplementary Fig. 4b, c**). To quantify the fold change in SspB membrane recruitment, we accounted for both the basal level of light-independent recruitment and the maximum light-induced membrane recruitment (**Fig. 2h**). SspB(nano) and SspB(micro) both displayed much larger fold changes in membrane recruitment than SspB(milli) with SspB(nano) slightly outperforming SspB(micro) across different light intensities (**Fig. 2i**). Comparing dissociation kinetics revealed that the dissociation half-times for SspB(nano, micro, milli) were relatively insensitive to the strength of the activating blue light although the three variants displayed different dissociation half-times that were correlated with their respective affinities (**Fig. 2j**). Based on its relatively low level of light-independent membrane recruitment and moderately large increase in light-dependent membrane localization, we determined SspB(micro) to be the most versatile for reconstituting membrane signaling reactions.

### Rapid and reversible Ras activation with spatial control

GTPases are toggled between their active, GTP-bound state and their inactive, GDP-bound state by GEFs and GAPs, respectively (**Fig. 1b**). To measure the kinetics of GAP- and GEF-mediated regulation of Ras, we visualized the change in membrane localization of a fluorescently labeled Ras(GTP) binding domain (RBD) derived from c-Raf kinase (**Fig. 3a**). Ras activation is often studied using GAPs and GEFs in isolation to drive reactions in a single irreversible direction. However, GAPs and GEFs can compete to establish distinct steady-state levels of Ras(GTP) ^28–30^. The time scale of Ras activation can also lead to different cell signaling fates ^31^. In many cases, receptor activation is followed by signal adaptation mechanisms that restore cellular Ras(GTP) concentrations to a basal level ^5^ (**Fig. 1c**). To mimic the dynamic rise and fall in Ras activity that is commonly observed in cell signaling, we established conditions to trigger the transient light-induced activation of Ras in the context of global inhibition imposed by the RasGAP NF1 (neurofibromin 1, subsequently referred to as RasGAP) (**Fig. 3b**). We conjugated purified Ras to SLBs in a physiologically relevant orientation using maleimide lipids, as previously described ^32^. To spatially control Ras activation, we designed and purified the catalytic domain of the Son of Sevenless homolog 1 (SOS1) RasGEF fused to SspB(micro) (subsequently referred to as GEF-SspB). Similar to SspB(micro) alone, we observed robust and transient recruitment of the GEF-SspB fusion to membrane-tethered iLID following patterned illumination with 488 nm light (**Fig. 3c, Supplemental Fig. 1a-b**, and **Supplemental Fig. 4d-h**). Immediately following light-induced membrane recruitment of GEF-SspB, we detected a local increase in AF546-RBD (active Ras) fluorescence intensity, indicating that we were successful in spatially controlling Ras activation (**Fig. 3c, d**). This local Ras(GTP) signal decayed back to baseline following membrane dissociation of GEF-SspB and RasGAP-mediated GTP hydrolysis (**Fig. 3b-d**). Importantly, the timescale of Ras activation and deactivation was comparable signaling dynamics observed in cells ^33^.

**Figure 3.**
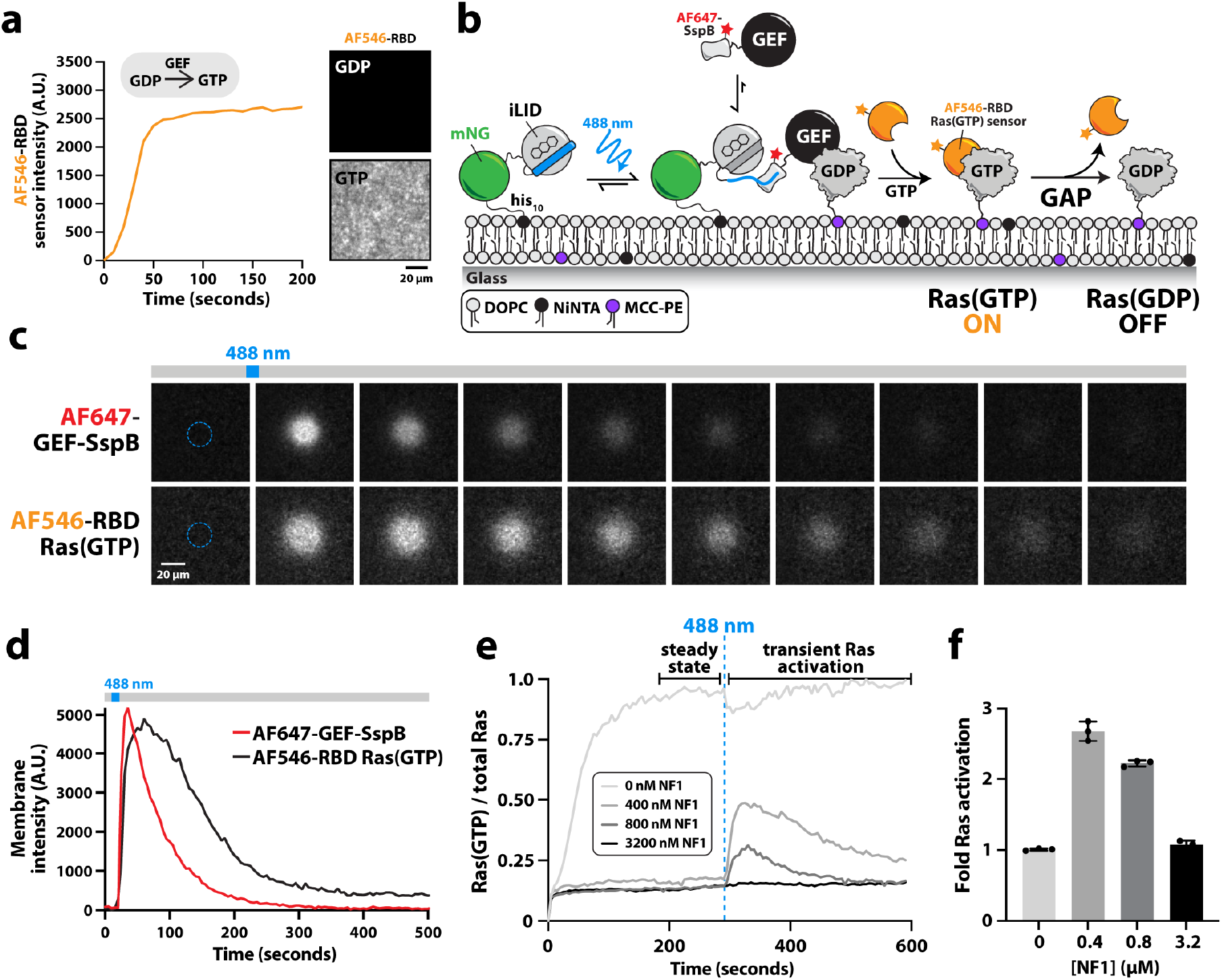
Reversible light-induced activation of membrane-tethered Ras GTPase. **(a)** Kinetics of membrane-anchored Ras activation in the presence of 50 nM RasGEF. Ras activation was monitored with 50 nM AF546-labeled Ras binding domain (RBD). Representative images show bulk membrane localization and specificity of AF546-RBD on bilayers with inactive Ras(GDP) and active Ras(GTP). **(b)** Cartoon schematic showing how light-induced activation of iLID drives membrane recruitment of AF647-GEF-SspB to locally activate membrane-anchored Ras. Global GAP inhibition reverses light-mediated activation of Ras. **(c)** Representative TIRF-M montages showing the time-dependent change in AF647-GEF-SspB(micro) membrane localization and Ras activation following local iLID activation with patterned 488 nm light (dashed blue circle). Images are separated by 25 seconds. **(d)** Kinetic traces showing membrane recruitment and dissociation of AF647-GEF-SspB(micro) and AF546-RBD following a 50 ms 488 nm light pulse. **(e)** Varying RasGAP concentrations (0-3200 nM) modulate the initial steady-state fraction of activated Ras and response to local light-induced activation by GEF-SspB(micro). **(f)** Quantification of the light-induced fold increase in Ras(GTP) as a function of global RasGAP concentration in **(e)**. Fold activation was calculated using (steady-state light-independent Ras activation + peak 488 nm light dependent activation)/steady-state light-independent activation. **(c-f)** Membrane composition: 92% DOPC, 2% NiNTA (+iLID bound), 2% MCC-PE (+Ras bound), and 4% PIP_2_ lipids. For all reactions, GEF refers to the catalytic domain of SOS1 (i.e. SosCat).

In vivo, the total RasGAP concentration can range from approximately 50 to 200 nM ^34,35^. However, these inhibitors are structurally variable and exist within complex regulatory networks making it difficult to estimate the total cellular concentration of “active” RasGAP. To determine the relationship between RasGAP concentration and steady-state Ras(GTP) levels, we tested our light-inducible Ras activation system over a range of RasGAP concentrations (**Fig. 3e, f**). For these experiments, we allowed reactions containing membrane-tethered Ras and GEF-SspB to reach an initial steady state of Ras activation in the presence of global GAP-mediated inhibition. Then, we used a transient pulse of 488 nm light guided by the DMD to drive membrane recruitment of GEF-SspB. We quantified the corresponding fold change in Ras(GTP) on membranes (**Fig. 3e, f**). In the absence of RasGAP, we observed no light-induced Ras activation because all the available Ras molecules were already GTP-bound due to unopposed global GEF activity. Conversely, when RasGAP was present at the high concentration of 3.2 µM, most Ras molecules remained GDP-bound and the change in Ras activation upon illumination was undetectable. In the intermediate RasGAP concentration range of 400-800 nM we measured robust increases in Ras(GTP) following light-induced activation with GEF-SspB. These Ras(GTP) signals decayed back to the baseline in minutes. We used RasGAP concentrations between 400 and 810 nM for all subsequent optogenetic experiments. By combining transient, light-mediated Ras activation and global RasGAP inhibition, we established a unique in vitro assay that resembles the spatiotemporal dynamics of Ras activation in vivo.

### Positive feedback drives Ras(GTP) amplification in the presence of global inhibition

In cells, GEFs bind activated GTPases or downstream products to generate recruitment-based feedback loops that amplify membrane signaling reactions ^4,36–38^. We hypothesized that GEF-mediated positive feedback would alter the spatiotemporal dynamics of light-inducible Ras activation in our reconstituted system. To test this, we designed a simple feedback (fb) circuit that relies on a minimal RasGEF catalytic domain derived from SOS1 fused to a Ras(GTP) binding domain (RBD) (**Fig. 4a**). This chimeric GEF, which we refer to as GEF(fb), binds with high specificity to Ras(GTP). Although SOS1 naturally exhibits positive feedback based on differential Ras binding to its catalytic and allosteric binding sites ^39^, our engineered positive feedback circuit allowed us to bypass complexity associated with SOS1 autoinhibition and receptor mediated activation. Measurement of Ras nucleotide exchange in the presence of GEF(fb) and the absence of GAP inhibition produced a sigmoidal kinetic trace consistent with strong recruitment-based positive feedback (**Fig. 4b**). Moreover, in the presence of GAP and high GEF(fb) concentrations (>30 nM), we occasionally observed bistable patterns of Ras(GDP) and Ras(GTP) on supported membranes (**Fig. 4c**). These patterns were reminiscent of previously reported phosphatidylinositol phosphate (PIP) lipid compositional patterns that form due to competing lipid kinases and phosphatases that exhibit positive feedback based on product binding ^20^. Depending on the balance of GAP and GEF activity, we could shift the system to steady-state conditions that either eliminated Ras(GTP) or caused Ras to be activated across the entire bilayer. This spontaneous symmetry breaking or bistability is an additional demonstration of GEF-mediated positive feedback in our system.

**Figure 4.**
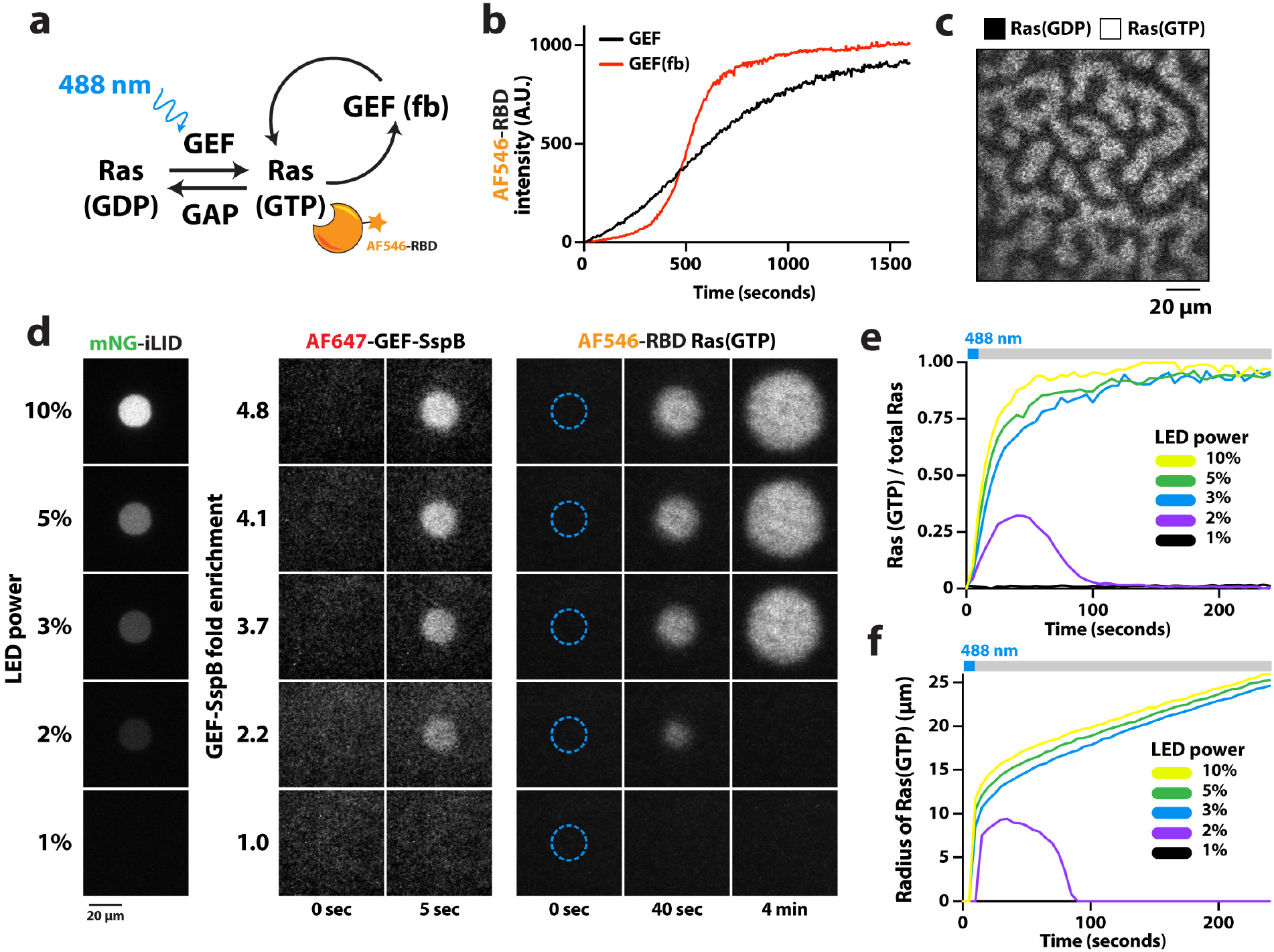
Light-induced Ras activation with positive feedback overcomes global GAP inhibition. **(a)** Diagram of light-inducible Ras activation module. Global RasGAP activity prevents Ras activation until light-induced recruitment of GEF-SspB generates a transient pool of Ras(GTP), which can be visualized via an AF546-RBD biosensor. This reaction is amplified by a positive feedback loop mediated by GEF(fb). **(b)** Hyperbolic Ras activation kinetics in the presence of 5 nM GEF and sigmoidal Ras activation kinetics in the presence of 1 nM GEF(fb). **(c)** Representative TIRF-M image showing Ras(GTP) (AF647-RBD) patterns spontaneously formed in the presence of GEF(fb) and RasGAP in the absence of 488 nm activating light. **(d)** Ras activation requires a threshold density of AF647-GEF-SspB to trigger GEF-mediated positive feedback. Representative TIRF-M images showing the region of iLID emission following excitation with patterned 488 nm light (left), AF647-GEF-SspB membrane localization with corresponding fold enrichment (middle), and the resulting Ras activation over time (right). **(e)** Change in the fraction of locally activated Ras over time for the experiment shown in **(d). (f)** Graph showing the change in the radii of the regions of AF546-RBD fluorescence over time for the experiment shown in **(d). (b)** Membrane composition: 98% DOPC and 2% MCC-PE (+Ras bound) lipids. **(c-f)** Membrane composition: 92% DOPC, 2% NiNTA (+iLID bound), 2% MCC-PE (+Ras bound), and 4% PIP_2_ lipids.

Membrane signaling reactions often require a threshold concentration of activator to overcome global inhibition ^40–43^. To determine the requirements to amplify and persistently outpace global GAP-mediated inhibition following local production of Ras(GTP), we varied the intensity of activating light to recruit a range of GEF-SspB (i.e. initiator) membrane densities (**Fig. 4d-f**). These experiments were performed under initial steady-state conditions that eliminated spontaneous Ras(GTP) pattern formation due to high RasGAP activity. Following a low, 2.2-fold membrane enrichment of GEF-SspB, we found that GEF(fb) in solution was unable amplify local Ras(GTP) enough to overcome global GAP inhibition (**Fig. 4d-f**). Instead, the local Ras(GTP) signal rapidly disappeared. Repetitive local stimulation with low-intensity light was similarly unable to trigger the Ras(GTP)-mediated positive feedback loop (**Supplementary Fig. 5a, b**). By contrast, when GEF-SspB membrane enrichment was greater than 3.7-fold, the Ras(GTP) signal not only persisted long after GEF-SspB dissociated but also resulted in the fluorescent region expanding across the bilayer (**Fig. 4d, f**). Our in vitro feedback system requires a threshold density of activated Ras molecules to overcome global inhibition and promote amplification through Ras(GTP)-mediated positive feedback.

To confirm that the spatial Ras(GTP) amplification we observed was indeed due to the presence of GEF-mediated positive feedback, we performed two identical Ras activation experiments with either GEF(fb) or an equivalent concentration of GEF lacking positive feedback (**Fig. 5a, b**). Whereas Ras activation following light-mediated activation (30% LED power) in the control experiment remained local and transient due to global GAP-mediated inhibition, the presence of the positive feedback loop allowed the Ras activation reaction to locally outcompete inhibition and resulted in an AF546-RBD (active Ras) signal that grew in intensity and expanded across the bilayer (**Fig. 5a**). The sharp boundaries and rapid expansion of the Ras(GTP) region we observed in the presence of GEF(fb) are reminiscent of a Fisher wave ^44^. This type of self-propagating wave arises when membrane diffusion and catalysis outpace global inhibition. Using the Fisher equation, we simulated the spatial distribution of Ras(GTP) molecules following light-induced Ras activation (**Fig. 5c, d**). This simulation depended on the kinetics of Ras activation (logistic growth), the membrane diffusion coefficient of Ras, and the reaction decay rate driven by GAP-mediated nucleotide hydrolysis. The resulting Fisher wave matched the spatial expansion we observed in our experiments with GEF-mediated positive feedback remarkably well. To confirm that this wave could not be attributed to simple diffusion of Ras(GTP) across the bilayer, we performed the same simulation with the logistic growth rate set to zero (**Fig. 5c, d**). We also ran simulations in the absence of both logistic growth and global inhibition (**Supplemental Fig. 6a-c**). As in our reconstitutions lacking positive feedback, the Ras(GTP) signal in the control simulations rapidly decayed due to membrane diffusion and global GAP-mediated inhibition following patterned activation of Ras.

**Figure 5.**
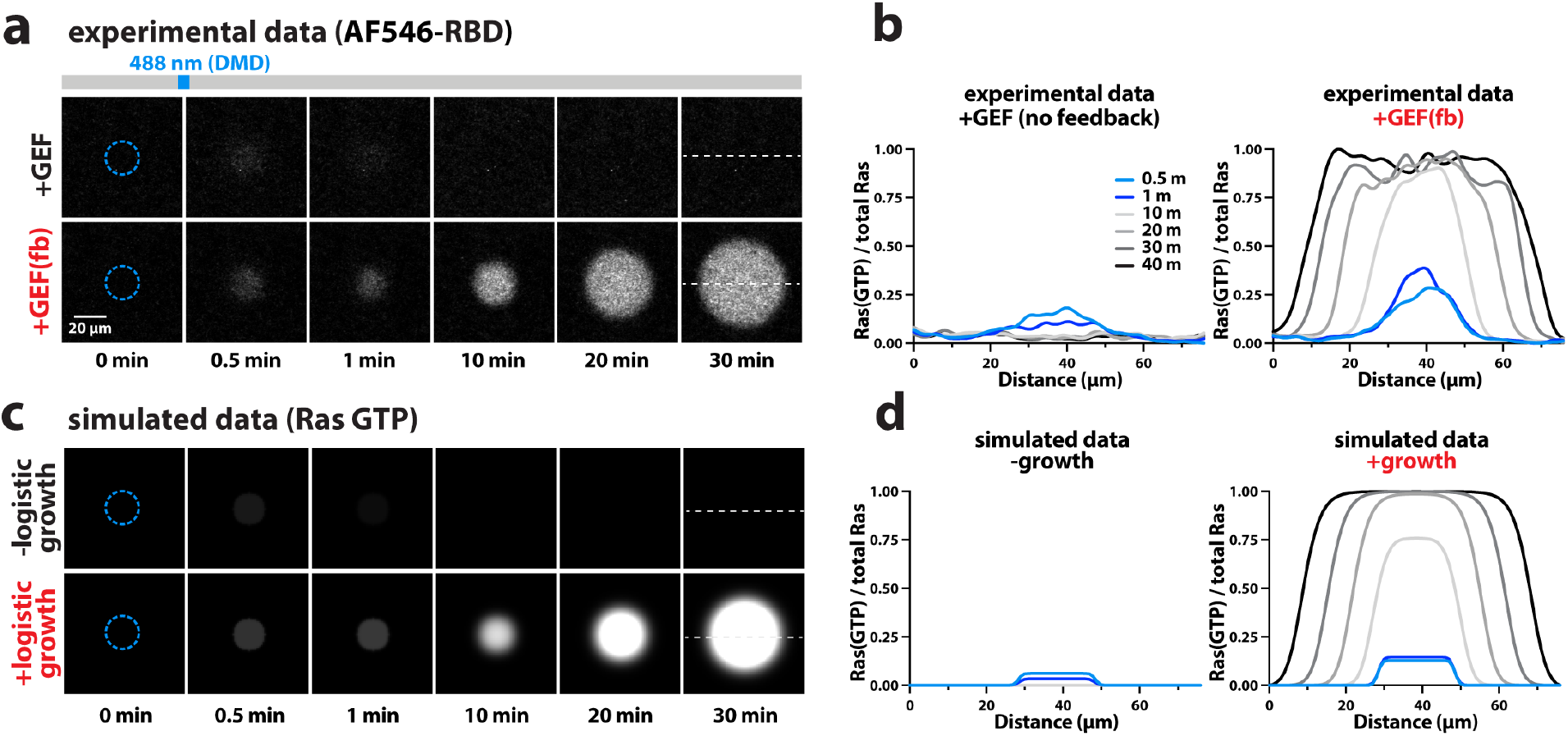
Positive feedback drives Ras(GTP) amplification and expansion as a Fisher wave. **(a)** Representative TIRF-M montages showing the time-dependent change in Ras activation following a 488 nm light pulse in a 20 µm-diameter circular membrane region in the presence of 10 nM GEF or 10 nM GEF(fb). **(b)** Intensity line scans showing the fraction of locally activated Ras over time for the experiments shown in **(a). (c)** Representative montages of simulated images obtained using the Fisher equation with and without a logistic growth term. The diffusion coefficient used was 1.12 µm^2^/sec which is the experimentally determined diffusion coefficient of membrane-tethered Ras. **(d)** Intensity line scans showing the fraction of locally activated Ras over time for the simulations shown in **(c). (a, b)** Membrane composition: 92% DOPC, 2% NiNTA (+iLID bound), 2% MCC-PE (+Ras bound), and 4% PIP_2_ lipids.

### Reconstitution of Ras(GTP)-PI3Kγ communication with global inhibition

Cell surface receptors and GTPase signaling are critical for the activation of class I phosphoinositide 3-kinases (PI3Ks), which catalyze PIP_2_ phosphorylation at the plasma membrane to generate PIP_3_^1,18^ . The PI3K gamma (PI3Kγ) variant is highly expressed in the immune system and is activated by GTP-bound Ras and GβGγ subunits (**Fig. 6a**) ^14–17,45^. We set out to determine how these factors localize and activate PI3Kγ using purified proteins on SLBs containing PIP_2_. We visualized kinase localization by imaging AF647-labeled PI3Kγ and simultaneously monitored lipid phosphorylation activity using a fluorescently labeled PIP_3_ biosensor derived from Btk (Bruton’s tyrosine kinase) (**Fig. 6b** and **Supplementary Fig. 7a**,**b**). We found that Ras(GTP) alone did not facilitate PI3Kγ membrane localization, whereas membrane-bound GβGγ could both recruit and activate PI3Kγ ^15^. Notably, when both Ras(GTP) and GβGγ were present, we observed a ∼2-fold increase in PI3Kγ membrane localization and a corresponding ∼2-fold increase in the rate of PIP_3_ generation compared to with GβGγ alone (**Fig. 6b** and **Supplementary Fig. 7b**). These results are consistent with the findings of previous biochemical studies of PI3Kγ activation ^15^. In cells, PTEN lipid phosphatase limits spurious PI3K signaling by converting PIP_3_ back to PIP_2_. By including PTEN in our reconstitution, we were able to balance GβGγ-dependent, Ras-independent generation of PIP_3_ by PI3Kγ. When we then rapidly activated a high density of Ras molecules by introducing a RasGEF to this reaction, we found that PI3Kγ activity was able to overcome PTEN-mediated inhibition (**Supplementary Fig. 7c**,**d**). PI3Kγ, membrane-bound Ras, membrane-bound GβGγ, and PTEN comprise a switchable, reconstituted signaling module that only leads to rapid PIP_3_ generation when Ras is activated by a GEF.

**Figure 6.**
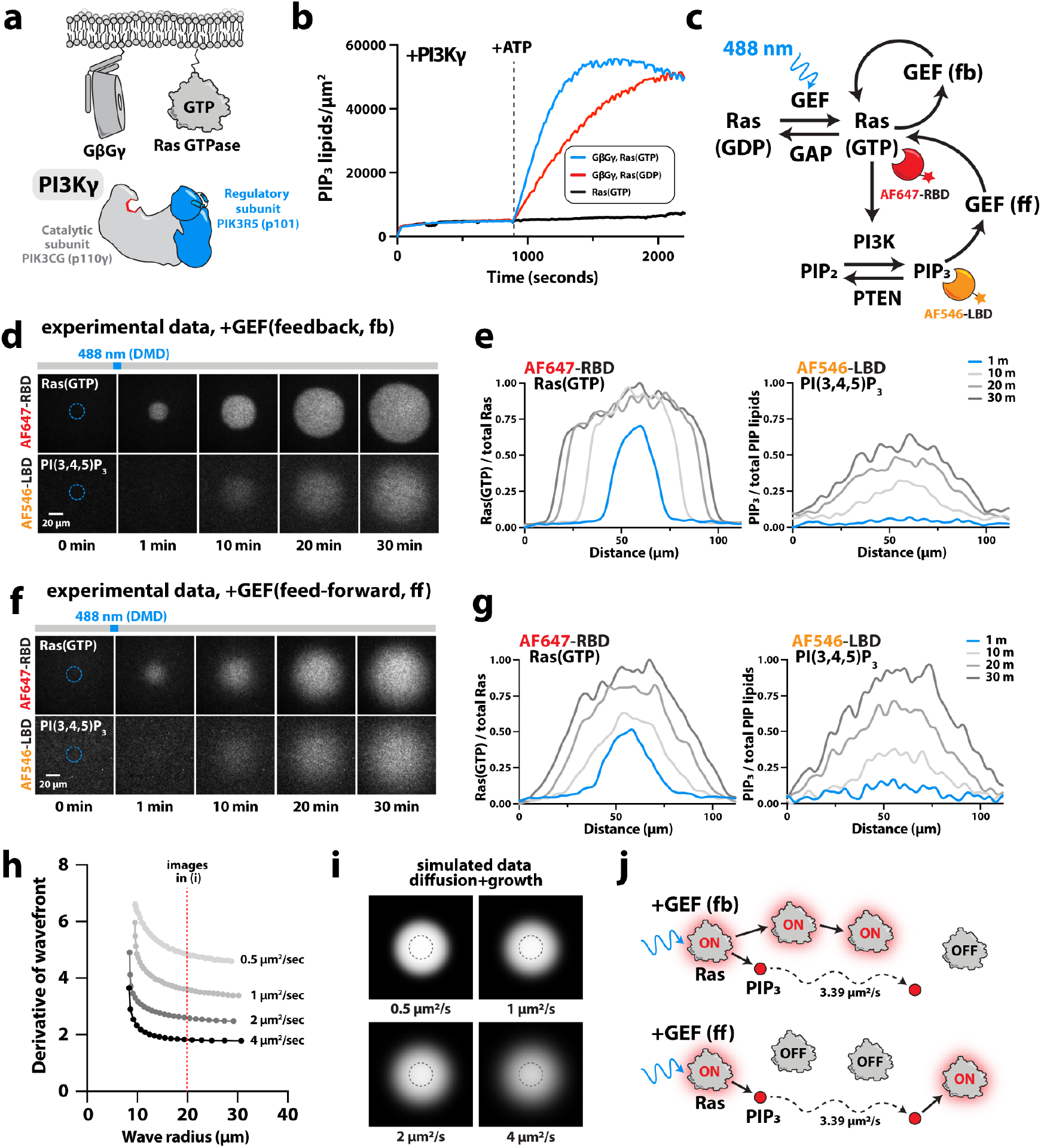
Diffusion modulates spatial coupling between Ras activation and PIP_3_ synthesis. **(a)** PI3Kγ is activated by membrane-tethered GβGγ and Ras(GTP). **(b)** Kinetics of PIP_3_ generation in the presence of Ras(GTP) and GβGγ. Reactions measured with 20 nM AF546-Btk. The dashed line marks the addition of 1 mM ATP. **(c)** Diagram of communication between light-induced Ras activation and PI3Kγ-dependent PIP_3_ generation. Adding GEF(fb) or GEF(ff) to the module introduces a positive feedback loop or a feed-forward loop, respectively. **(d)** Representative montages showing the time-dependent change in Ras activation and PIP_3_ generation following a 488 nm light pulse (blue circle), in the presence of the GEF(fb)-mediated positive feedback loop. **(e)** Intensity line scans corresponding to the experiment shown in **(d). (f)** Representative montages showing the time-dependent change in Ras activation and PIP_3_ generation following a 488 nm light pulse (blue circle), in the presence of the GEF(ff)-mediated feed-forward loop. **(g)** Intensity line scans corresponding to the experiment shown in **(f). (h)** Ras(GTP) Fisher waves were simulated with varying Ras membrane diffusivity. The steepness (i.e. derivative) of the Ras(GTP) wave edge changes based on the diffusivity of Ras as the activated region expands. **(i)** Images of simulated Ras(GTP) Fisher waves following expansion to a radius of 20 µm. The dashed circle represents the initial region of Ras activation. **(j)** Model for different wave characteristics observed in the presence of GEF(fb) and GEF(ff). **(d-g)** Membrane composition: 92% DOPC, 2% NiNTA (+iLID bound), 2% MCC-PE (+Ras bound), and 4% PIP_2_ lipids, +GβGγ-farnesyl.

### Diffusion modulates spatial coupling between Ras(GTP) and PIP_3_ synthesis

To decipher the spatial coupling between Ras(GTP) and PIP_3_ production on membranes, we combined our reconstitution of the Ras-PI3Kγ module in the presence of global inhibitors (i.e. RasGAP and PTEN) with iLID-SspB-dependent light-inducible Ras activation (**Fig. 6c**). First, we examined the spatiotemporal amplification of Ras(GTP) and PIP_3_ following transient membrane recruitment of the GEF-SspB initiator in the presence of GEF(fb)-mediated positive feedback (**Fig. 6c-e**). As previously observed, light-induced recruitment of GEF-SspB resulted in the local production of Ras(GTP), which was then spatially amplified by GEF(fb). Under these conditions, we found that local PI3Kγ activity was also able to outcompete PTEN-mediated dephosphorylation of PIP_3_. Although PIP_3_ generation lagged behind Ras(GTP), the region of AF546-LBD biosensor fluorescence tracked the Ras(GTP) signal and expanded beyond the original light-activated region (**Fig. 6d, e**). These results demonstrate communication between light-inducible Ras activation and PI3Kγ signaling. A minimal GEF-mediated positive feedback loop is sufficient for both reactions to overcome global GAP- and PTEN-mediated inhibition.

In vivo, GEFs can mediate feed-forward loops that drive GTPase activation based on PIP_3_-dependent membrane recruitment and activation ^4,36^. Having successfully reconstituted Ras(GTP)-PI3Kγ communication in the presence of GEF-mediated positive feedback, we aimed to reconstitute a GEF-mediated feed-forward (ff) circuit that could amplify Ras downstream of PI3Kγ-mediated PIP_3_ production (**Fig. 6c**). Much like the GEF(fb)-mediated feedback loop, we engineered a feed-forward loop by fusing a Btk-derived PIP_3_ lipid-binding domain (LBD) to a SOS1 GEF domain to generate a chimera we refer to as GEF(ff). Following transient, light-induced membrane recruitment of the GEF-SspB initiator, GEF(ff) was indeed able to drive the spatial expansion of Ras(GTP) and PIP_3_ generation on membranes (**Fig. 6f, g**). These amplification patterns exhibited similar dynamics to those we observed previously in the presence of GEF(fb). The results of introducing GEF(fb) and GEF(ff) to our reconstituted pathway provide molecular evidence for the sufficiency of simple, GEF-mediated feedback circuits to produce spatiotemporal PIP_3_ signal amplification, an essential step in numerous cell signaling pathways.

Although the PIP_3_ signal was spatially correlated with Ras(GTP) in the presence of both GEF(fb) and GEF(ff), we observed a pronounced difference in the boundary steepness of Ras(GTP) waves when comparing the two feedback circuits (**Fig. 6d-g**). Positive feedback mediated by GEF(fb) invariably resulted in a uniformly bright Ras(GTP) region with a steep wavefront (**Fig. 6d, e**). By contrast, the PIP_3_ wavefronts were less defined than those of Ras(GTP). Importantly, when Ras activation depended on the generation of rapidly diffusing PIP_3_ lipids due to GEF(ff), the Ras(GTP) wavefront more closely resembled the gradual boundary of the PIP_3_ region. Overall, these observations are consistent with the markedly different diffusion coefficients of membrane-tethered Ras (∼0.5 µm^2^/sec) and lipids within the supported membrane (∼3 µm^2^/sec) (**Supplementary Fig. 8**). To model the effect of diffusivity on wavefront expansion and steepness, we simulated Fisher wave growth using a range of diffusion coefficients for the membrane bound species (i.e. Ras or PIP_3_) (**Fig. 6h, i**). Following spatially patterned activation of Ras(GTP), simulated Fisher waves expanded at rates proportional to the diffusion coefficient. However, the steepness of the wave boundary decreased as diffusivity increased (**Fig. 6h**), a result that is consistent with the spatial coupling observed between Ras(GTP) and PIP_3_ in our reconstitution. These results support a model in which the lateral membrane diffusion of reacting species together with regulatory circuit architecture modulate the spatial coupling and evolution of membrane signaling reactions (**Fig. 6j**).

## DISCUSSION

This study establishes a new framework for reconstituting communication and feedback between membrane signaling reactions involving small GTPases, PIP lipids, and the enzymes that regulate their production. We devised an approach by which we can observe unique steady-state conditions in the presence of both activators and inhibitors of Ras GTPase. By suppressing spurious activation of Ras through global, GAP-mediated inhibition we mimic cellular membrane signaling states prior to receptor agonist-induced pathway activation. We then utilize transient, light-induced membrane recruitment of a RasGEF via the iLID-SspB system to trigger the local activation of membrane-anchored Ras GTPase. We demonstrate that reactions at steady state can be rapidly shifted into new signaling regimes in a spatially controlled manner. Importantly, the light-based signaling inputs and corresponding responses occur on a biologically relevant time scale (i.e. seconds to minutes). Using this framework, we examine two membrane recruitment-based positive feedback circuits and the requirements for a reconstituted Ras-PI3Kγ signaling module to overcome global inhibition and amplify in space and time. We demonstrate that an initial threshold density of membrane-localized GEF is required to transiently outcompete GAP-mediated Ras inhibition. This triggers feedback-dependent signal amplification in the form of an expanding Fisher wave. When the threshold for Ras activation is not achieved, the membrane signaling reaction resets to the “OFF” state with dynamics reminiscent of cellular signal adaptation. For a feed-forward circuit that links Ras activation to PIP_3_ production, the spatiotemporal coupling of the two reactions is regulated by differences in lateral membrane diffusion of Ras(GTP) and PIP_3_ lipids. Our observations support previous stochastic kinetic modeling showing that molecules with slower diffusion coefficients exhibit more robust polarity in two-component reaction-diffusion systems with positive feedback based on product binding ^7^.

In vivo, communication between Ras and PIP_3_ is established through an extensive network of positive and negative feedback loops that are challenging to decipher due to redundancy, network crosstalk, and signal adaptation. Positive feedback within the Ras-PI3K signaling pathway can at least in part be attributed to GEFs that are recruited to the plasma membrane by active Ras itself and direct interactions with downstream PIP_3_. These GEFs function analogously to the chimeric GEFs used in our assay although they contain additional layers of regulation, such as reversible autoinhibition. In nature, processive membrane association of RasGEFs also plays a role in increasing local Ras(GTP) density and promiscuous GEFs that bind multiple classes of small GTPases via their allosteric or catalytic domains amplify signaling events beyond direct Ras signaling ^21,32,46,47^. There is also a large body of evidence supporting Ras-independent feedback circuits mediated by downstream effectors in the PI3K signaling pathway. For example, PIP_3_ generation promotes the recruitment of GEFs such as Vav, P-Rex, and the DOCK/ELMO complex, which activate Rac GTPase ^36,48–52^. GTPases including Rac, Cdc42, and RhoG, which are activated in parallel and downstream of Ras, can also stimulate PI3K activity. PI3K-mediated PIP_3_ generation and Rac activation lead to the formation of branched actin networks which deform the membrane and produce force at the leading edge of migratory cells ^53,54^. Dendritic actin networks also promote PIP_3_ production by delivering signaling molecules including PI3K and various GEFs to the cell cortex in a polarized manner ^12,55–59^. Cell biological evidence points to the presence of myriad positive feedback circuits in signaling networks and computational modeling has led to speculation about considerably more. Few of these circuits have been interrogated through biochemical reconstitution. Our study represents an exploration of a single regulatory node that is known to modulate hundreds of signaling molecules in living cells.

Small GTPases are understood to be key activators of class I PI3Ks and the relationship between these classes of molecules is at the core of numerous signaling cascades ^47,57^. Best known for its role in myeloid cells, the PI3Kγ isoform has been shown to be activated by GPCR-derived GβGγ subunits and GTP-bound Ras GTPases in vivo and in vitro ^14,16,17,45^. In this study, we directly observed PIP_3_ generation by PI3Kγ in the presence of membrane-bound Ras and GβGγ. Using light-induced membrane recruitment of a GEF to initiate the local activation of Ras, we reconstitute communication between this GTPase and PI3Kγ-mediated production of PIP_3_ lipids. In agreement with previous biochemical studies ^15^, we demonstrate that Ras(GTP) alone is unable to activate PI3Kγ. However, the presence of both Ras(GTP) and GβGγ leads to enhanced PI3Kγ membrane localization and PIP_3_ generation. In the presence of global PTEN phosphatase activity, we found that Ras activation is required for PI3Kγ-dependent lipid phosphorylation.

Cells establish precise, system-level steady-state signaling conditions by balancing complex networks of activators and inhibitors. The total concentrations of cell surface receptors, small GTPases, GEFs, GAPs, lipid kinases, and lipid phosphatases vary between cell types, creating unique thresholds for activation, rates of signal amplification, and cellular outcomes ^60^. The total cellular concentration of RasGAPs and RasGEFs can range from 50 to 200 nM ^34,35^. However, the system-level concentrations and activities of GAPs and GEFs are difficult to estimate due to our incomplete understanding of their regulatory mechanisms (e.g. autoinhibition). This makes it challenging to approximate biologically relevant strengths of global Ras activation and inhibition. In our assay, we use a single RasGAP (NF1) at concentrations between 400 and 800 nM which we experimentally determined to be sufficient to suppress most GEF-mediated Ras activation. This global, GAP-driven inhibition stands in for the intricate, system-level Ras regulatory modules that exist across cell types.

Steady-state cellular conditions can also be altered by mutations to the molecules that comprise signaling networks. Dysregulation of key nodes like Ras and PI3K has long been known to have severe cellular consequences which has flagged these molecules and their associated regulators as important cancer therapeutic targets ^2,13,18,19,29,43,61,62^. The inhibitors we used in our in vitro assay, NF1 RasGAP and PTEN phosphatase, are well-characterized tumor suppressors that prevent aberrant Ras and PIP_3_ signaling ^63,64^. Cancer cell lines which have lost NF1 RasGAP expression exhibit elevated steady-state levels of Ras(GTP) ^65^. Conversely, enhancing the membrane binding of PTEN in cells expressing constitutively active PI3K leads to a marked decrease in membrane PIP_3_^66^ . Biochemical reconstitution of GTPase and PIP lipid signaling in the presence of mutated GTPases, kinases, and their respective regulators will shed light on how the relationships between these molecules determine the properties of healthy and diseased cells. In addition to influencing steady-state conditions, alterations to regulatory modules via mutations or relative protein concentrations can affect how signaling networks transition between steady states. For this reason, it is important to monitor the dynamic evolution of reactions in addition to measuring steady state conditions when investigating perturbations to signaling networks.

One aspect of cell signaling that is not explored in this study is delayed negative feedback, which can be broadly categorized as long-range or local. Long-range negative feedback mediated by cortical membrane-cytoskeletal tension allows for the establishment and maintenance of asymmetry over the entire length of a polarized cell ^67,68^. By contrast, local negative feedback relies on the products of specific biochemical reactions to recruit inhibitors and is fundamental for signal adaptation. In the context of directed cell migration, local negative feedback prevents cells from becoming “polarity-locked”, thus maintaining their flexibility and sensitivity to changes in environmental signals ^3,5,9,48,69,70^. GAPs, which directly oppose GEF activity, are natural candidates for mediating local delayed negative feedback ^59,71^. Unsurprisingly, various GAPs have been shown to serve functions in directed cell migration ^72,73^ and signal adaptation ^48^. Like GEFs, some GAPs may be directly recruited to the membrane by small GTPases ^73^. Additionally, there is evidence for different classes of GAPs binding to PIP_3_ and a downstream product, PI(3,4)P_2_^74–77^ . PI(3,4)P_2_ which is converted from PIP_3_ by 5-phosphatases has also been shown to play a role in adaptive endocytic pathways through which activated receptors are turned over at the leading edge of migrating cells ^78^. Interestingly, actin networks, which promote PI3K activity and mediate long-range negative feedback, also participate in local negative feedback by inhibiting Rac and Rho GTPases ^48,79,80^. Filamentous actin is hypothesized to deliver GAPs directly to the membrane and may also physically drive the dissociation of signaling molecules through network treadmilling ^59,81^. Although we have relied on transient inputs and global inhibition to build reversibility into our assay, it is feasible to design similar assays to investigate the principles of adaptive signaling modules in the presence of a sustained light inputs in the future.

## Supporting information

Supplementary File_FisherWave simulation

## ACKNOWLEDGMENTS

Research was supported by the University of Oregon Start-up funds (S.D.H.), National Science Foundation CAREER Award (S.D.H., MCB-2048060), and the University of Oregon Presidential Undergraduate Research Scholars (PURS) program (A.O.C.). We thank Peter Bieling (King’s College, London UK) for critical feedback or our manuscript draft. We thank Doug Tischer and Orion Weiner (University of California at San Francisco) for providing plasmids containing genes that encode iLID and SspB. We thank Sofia Carlson (University of Oregon) for assisting with preliminary experiments involving iLID-SspB membrane recruitment. We thank Tristan Ursell and Raghuveer Parthasarathy (University of Oregon, Department of Physics) for discussions about Fisher wave simulations and analysis of reaction diffusion systems. The content is solely the responsibility of the authors and does not necessarily represent the official views of the National Science Foundation.

## SUPPORTING INFORMATION

Supplementary Figures S1-S8

## AUTHOR CONTRIBUTIONS

Resources: S.D.H., A.O.C., S.D.

Experiments and investigation: S.D.H., A.O.C., S.D.

Data Analysis: S.D.H., A.O.C., S.D.

Conceptualization: S.D.H., S.D.

Interpretation: S.D.H., A.O.C., S.D.

Data curation: S.D.H., A.O.C., S.D.

Writing – Review and editing: S.D.H., A.O.C., S.D.

Writing – Original draft: S.D.H., S.D.

Supervision: S.D.H.

Project administration: S.D.H.

Funding acquisition: S.D.H.

## DATA AVAILABILITY

All the information needed for interpretation of the data is presented in the manuscript or the supplementary material. Plasmids related to this work are available upon request.

## MATERIALS & METHODS

### Molecular Biology

The following genes were used as templates for PCR to clone plasmids used for recombinant protein expression: *PIK3CG* (mouse 1-1102aa; Uniprot Accession #Q9JHG7), *PIK3R5* (mouse 1-871aa; Uniprot Accession #Q5SW28), *BTK* (bovine 1-659aa; Uniprot Accession #Q3ZC95), *GNB1/GBB1* (Gβ_1_, bovine 1-340aa; Uniprot Accession #P62871), *GNG2/GBG2* (Gγ_2_, bovine 1-71aa; Uniprot Accession #P63212). The following plasmids were purchased as cDNA clones from Horizon (PerkinElmer), formerly known as Open Biosystems and Dharmacon: mouse *PIK3CG* (clone #BC051246, Cat #MMM1013-202770664) and mouse *PIK3R5* (clone #BC128076, cat #MMM1013-211693360), human *CYTH3*/Grp1 (clone #4811560, cat #MHS6278-202806616). Genes encoding bovine Gβ_1_ and Gγ_2_ were derived from the following plasmids: YFP-Gβ_1_ (Addgene plasmid #36397) and YFP-Gγ_2_ (Addgene plasmid #36102). These Gβ_1_ and Gγ_2_ containing plasmids were kindly provided to Addgene by Narasimhan Gautam^82^. In this study, we used a previously described mutant form of Btk with mutations in the peripheral PIP_3_ binding site (R49S/K52S) ^83,84^. The Btk peripheral site mutant was PCR amplified using a plasmid provided by Jean Chung (Colorado State, Fort Collins) that contained the following coding sequence: his6-SUMO-Btk(PH-TH, R49S/K52S)-EGFP. Refer to supplementary text to see exact peptide sequence of every protein purified in this study. For cloning, genes were PCR amplified using AccuPrime *Pfx* master mix (ThermoFisher, Cat #12344040) and iProof High-Fidelity master mix (Bio-Rad, Cat #172-5320) combined with a restriction digested plasmids using Gibson assembly^85^. Refer to the Supplementary Information for a complete list of plasmids used in this study. Information about the specific peptide sequences for recombinantly expressed and purified proteins is organized in the Supplementary Information document. The complete open reading frames of all vectors used in this study were sequenced to ensure the plasmids lacked deleterious mutations. Refer to the plasmid inventory in the Supplementary Information for the complete list of protein expression vectors used in this study.

### Protein purification

#### mNeonGreen-iLID

Gene sequence encoding the iLID was expressed in Rosetta 2 pLysS *E. coli* as a his_10_-mNG-iLID fusion protein. Bacteria were grown at 37°C in Terrific Broth until it reached OD_600_ = 0.8. Cultures were shifted to 18°C for 1 hour then induced with 0.05 mM IPTG. Expression was allowed to continue for 22 hours before harvesting. Cells were lysed by sonication in buffering containing 50 mM Na_2_HPO_4_ [pH 8.0], 400 mM NaCl, 0.4 mM BME, 1 mM PMSF, 100 μg/mL DNase. Lysate was centrifuged at 15,000 rpm (35000 x g) for 60 minutes in a Beckman JA-20 rotor at 4ºC. Lysate was circulated over a 5 mL HiTrap chelating column (Cytiva, Cat# GE17-0409-01) inoculated with 100 mM CoCl_2_ for 10-15 minutes. Column was washed with 50 mM Na_2_HPO_4_ [pH 8.0], 400 mM NaCl, 10 mM imidazole, 0.4 mM BME. Bound protein was eluted in 50 mM Na_2_HPO_4_ [pH 8.0], 400 mM NaCl, 500 mM imidazole, 0.4 mM BME. Peak fractions were pooled and dialyzed against 4 liters of buffer containing 25 mM Na_2_HPO_4_ [pH 8.0], 400 mM NaCl, 500 mM imidazole, 0.4 mM BME. Dialysate was recirculated over a 5 mL HiTrap chelating column and concentrated in a 5 kDa MWCO Vivaspin 20 concentrator to a volume of 5 mL before loading onto a 124 mL Superdex 75 column equilibrated in 20 mM HEPES [pH 7.0], 200 mM NaCl, 10% glycerol, 1 mM TCEP. Peak fractions were pooled, concentrated to a minimum concentration of 200 µM, and snap frozen in liquid nitrogen for storage at -80ºC. The his10 tag caused the his10-mNG-iLID protein aggregate after long-term storage in the -80ºC. Before performing SLB experiments, we removed his10-mNG-iLID aggregated by size exclusion chromatography (i.e. Superdex 75).

#### SspB(milli, micro, nano)

Genes encoding SspB were expressed as his_10_-SUMO3-KCK-SpyCatcher-SspB fusion proteins in Rosetta 2 pLysS *E. coli*. Note that the SpyCatcher was simply used as a generic place holder fusion protein. It was not combined with SpyTag in this study. Bacteria were grown at 37°C in Terrific Broth to OD_600_ = 0.8. Cultures were then shifted to 18°C for 1 hour before inducing with 0.05 mM IPTG. Expression was allowed to continue for 22 hours before harvesting. Cells were lysed in 50 mM Na_2_HPO_4_ [pH 8.0], 400 mM NaCl, 0.4 mM BME, 1 mM PMSF, 100 μg/mL DNase by sonication. Lysate was centrifuged at 15,000 rpm (35000 x g) for 60 minutes in an Avanti JA-17 rotor at 4ºC. Lysate was circulated over a 5 mL HiTrap chelating column (Cytiva, Cat# GE17-0409-01) inoculated with 100mM CoCl_2_ for ten minutes. Column was washed with 50 mM Na_2_HPO_4_ [pH 8.0], 400 mM NaCl, 10 mM imidazole, 0.4 mM BME. Bound protein was eluted in 50 mM Na_2_HPO_4_ [pH 8.0], 400 mM NaCl, 500 mM imidazole, 0.4 mM BME. Peak fractions were pooled and his_6_-SenP2 (SUMO protease) was added to cleave off the his_10_-SUMO3 tag. The solution was dialyzed overnight against 4 liters of buffer containing 20 mM HEPES [pH 7.0], 200 mM NaCl, 10% glycerol, 0.4 mM BME. Dialysate was concentrated in a 5 kDa MWCO Vivaspin 20 concentrator to a volume of 5 mL before loading onto a 124 mL Superdex 75 column equilibrated in 20 mM HEPES [pH 7.0], 200 mM NaCl, 10% glycerol, 1 mM TCEP. Peak fractions were pooled and snap frozen in liquid nitrogen for storage at -80ºC.

#### GEF-SspB(milli, micro, nano)

Gene sequences encoding his_10_-SUMO3-KCK-SosCat(564-1049aa)-SspB were expressed in Rosetta 2 pLysS *E. coli* to generate chimeric light regulated GEFs. Bacteria were grown at 37°C in Terrific Broth until it reached OD_600_ = 0.8. Cultures were shifted to 18°C for 1 hour then induced with 0.05 mM IPTG. Expression was allowed to continue for 22 hours before harvesting. Cells were lysed in 50 mM Na_2_HPO_4_ [pH 8.0], 400 mM NaCl, 0.4 mM BME, 1 mM PMSF, 100 μg/mL DNase by sonication. Lysate was centrifuged at 15,000 rpm (35000 x g) for 60 minutes in a Beckman JA-20 rotor at 4ºC. Lysate was circulated over a 5 mL HiTrap chelating column (Cytiva GE17-0409-01) inoculated with 100mM CoCl_2_ for ten minutes. Column was washed with 50 mM Na_2_HPO_4_ [pH 8.0], 400 mM NaCl, 10 mM imidazole, 0.4 mM BME. Bound protein was eluted in 50 mM Na_2_HPO_4_ [pH 8.0], 400 mM NaCl, 500 mM imidazole, 0.4 mM BME. Peak fractions were pooled and SenP2 protease was added to cleave off the his_10_-SUMO3 tag. The solution was dialyzed overnight against 4 liters of buffer containing 25 mM Na_2_HPO_4_ [pH 8.0], 400 mM NaCl, 500 mM imidazole, 0.4 mM BME. Dialysate was recirculated over a 5 mL HiTrap chelating column and concentrated in a 5 kDa MWCO Vivaspin 20 concentrator to a volume of 5 mL before loading onto a 124 mL Superdex 75 column equilibrated in 20 mM HEPES [pH 7.0], 200 mM NaCl, 10% glycerol, 1 mM TCEP. Peak fractions were pooled, concentrated to a minimum concentration of 200 µM, and snap frozen in liquid nitrogen for storage at -80ºC.

#### H-Ras

A gene sequence encoding human H-Ras was expressed in BL21(DE3) Rosetta *E. coli* as a his_6_-TEV-H-Ras(1-181aa, C118S) fusion as previously described ^86^. Bacteria were grown at 37°C in Terrific Broth until it reached OD_600_ = 0.8. Cultures were shifted to 18°C for 1 hour then induced with 0.1 mM IPTG. Expression was allowed to continue for 20 hours before harvesting. Cells were lysed in 50 mM Na_2_HPO_4_ [pH 8.0], 300 mM NaCl, 10 mM imidazole, 4 mM BME, 1 mM PMSF, 100 μg/mL DNase by sonication. Lysate was centrifuged at 15,000 rpm (35000 x *g*) for 60 minutes in a Beckman JA-20 rotor at 4ºC. NiNTA resin was added to the supernatant and allowed to bind for 1 hour, then washed with 50 mM Na_2_HPO_4_ [pH 8.0], 300 mM NaCl, 10 mM imidazole, 4 mM BME. Bound protein was eluted in 50 mM Na_2_HPO_4_ [pH 8.0], 400 mM NaCl, 500 mM imidazole, 4 mM BME. Peak fractions were pooled and a TEV protease was added to cleave off the his_6_-TEV tag. The solution was dialyzed against 4 liters of buffer containing 50 mM Na_2_HPO_4_ [pH 8.0], 300 mM NaCl, 5% glycerol, 0.4 mM BME. Dialysate containing cleaved protein was recirculated over NiNTA resin. Flowthrough was concentrated in a 10 kDa MWCO Amicon-Ultra concentrator before loading onto a 124 mL Superdex 75 column equilibrated in 20mM Tris [pH 8.0], 200 mM NaCl, 10% glycerol, 0.5 mM TCEP. Peak fractions were pooled, concentrated to a minimum concentration of 500 µM, and snap frozen in liquid nitrogen for storage at -80ºC.

#### SNAP-RBD

A gene sequence encoding the Ras binding domain (RBD) derived by human cRaf kinase was expressed in Rosetta 2 pLysS *E. coli* cells as a his_6_-GST-PP-SNAP-GSGSGS-RBD (Raf1, K65E) fusion protein as previously described^87^. Bacteria were grown at 37°C in Terrific Broth until it reached OD_600_ = 0.8. Cultures were shifted to 18°C for 1 hour then induced with 0.5 mM IPTG. Expression was allowed to continue for 22 hours before harvesting. Cells were lysed in 50 mM Na_2_HPO_4_ [pH 8.0], 400 mM NaCl, 0.4 mM BME, 1 mM PMSF, 100 μg/mL DNase, 1 mg/ml lysozyme by sonication. Lysate was centrifuged at 15,000 rpm (35000 x g) for 60 minutes in a Beckman JA-20 rotor at 4ºC. Lysate was circulated over a 5 mL HiTrap chelating column (Cytiva, Cat# GE17-0409-01) inoculated with 100mM CoCl_2_ for ten minutes. Column was washed with 50 mM Na_2_HPO_4_ [pH 8.0], 400 mM NaCl, 10 mM imidazole, 0.4 mM BME. Bound protein was eluted in 50 mM Na_2_HPO_4_ [pH 8.0], 400 mM NaCl, 500 mM imidazole, 0.4 mM BME. Peak fractions were pooled and the his_6_-GST tag was cleaved off with PreScission protease (derived from human rhinovirus 3C protease), while dialyzing into buffer containing 20 mM HEPES [pH 7.5], 400 mM NaCl, 5% glycerol, 0.4 mM BME. Dialysate containing cleaved protein was recirculated over a 5 mL HiTrap chelating column. Flowthrough was concentrated in a 5 kDa MWCO Vivaspin 20 concentrator to a volume of 5 mL before loading onto a 124 mL Superdex 75 column equilibrated in 20mM HEPES [pH 7.0], 200 mM NaCl, 10% glycerol, 1 mM TCEP. Peak fractions were pooled and snap frozen in liquid nitrogen for storage at -80ºC.

#### GEF

A gene sequence encoding the catalytic domain of SOS1 was expressed in BL21 Trigger/pRIL *E. coli* cells as a his_6_-TEV-SosCat(564-1049aa) fusion protein as previously described ^39^. This protein is referred to as “GEF” throughout the manuscript. Bacteria were grown at 37°C in Terrific Broth to OD_600_ = 0.8. Cultures were shifted to 18°C for 1 hour then induced with 0.5 mM IPTG. Expression was allowed to continue for 22 hours before harvesting. Cells were lysed in 50 mM Na_2_HPO_4_ [pH 8.0], 400 mM NaCl, 10 mM imidazole, 4 mM BME, 1 mM PMSF, 100 μg/mL DNase by sonication. Lysate was centrifuged at 15,000 rpm (35000 x g) for 60 minutes in a Beckman JA-20 rotor at 4ºC. NiNTA resin was added to the supernatant and allowed to bind for 1 hour, then washed with 50 mM Na_2_HPO_4_ [pH 8.0], 400 mM NaCl, 10 mM imidazole, 4 mM BME. Bound protein was eluted in 50 mM Na_2_HPO_4_ [pH 8.0], 400 mM NaCl, 500 mM imidazole, 4 mM BME. Peak fractions were pooled and a TEV protease was added to cleave off the his_6_-TEV tag. The solution was dialyzed against 4 liters of buffer containing 20 mM HEPES [pH 7.0], 400 mM NaCl, 20 mM imidazole, 5% glycerol 0.4 mM BME. Dialysate containing cleaved protein was recirculated over NiNTA resin. Flowthrough was concentrated in a 30 kDa MWCO Amicon-Ultra concentrator to a volume of 5 mL before loading onto a 124 mL Superdex 75 column equilibrated in 20 mM Tris [pH 8.0], 150 mM NaCl, 10% glycerol, 1 mM TCEP. Peak fractions were pooled and snap frozen in liquid nitrogen for storage at -80ºC.

#### RasGAP

The RasGAP domain derived from Neurofibromatosis type 1 (NF1) was expressed in Rosetta 2 pLysS *E. coli* as a his6-MBP-TEV-NF1(1-333aa) fusion. Bacteria were grown at 37°C in Terrific Broth until it reached OD_600_ = 0.8. Cultures were shifted to 18°C for 1 hour then induced with 0.5 mM IPTG. Expression was allowed to continue for 22 hours before harvesting. Cells were lysed in 50 mM Na_2_HPO_4_ [pH 8.0], 400 mM NaCl, 0.4 mM BME, 1 mM PMSF, 100 μg/mL DNase, 1 mg/ml lysozyme by sonication. Lysate was centrifuged at 15,000 rpm (35000 x g) for 60 minutes in a Beckman JA-20 rotor at 4ºC. Lysate was circulated over a 5 mL HiTrap chelating column (Cytiva, Cat# GE17-0409-01) inoculated with 100 mM CoCl_2_ for ten minutes. Column was washed with 50 mM Na_2_HPO_4_ [pH 8.0], 400 mM NaCl, 10 mM imidazole, 0.4 mM BME. Bound protein was eluted in 50 mM Na_2_HPO_4_ [pH 8.0], 400 mM NaCl, 500 mM imidazole, 0.4 mM BME. Peak fractions containing the eluted protein were pooled, mixed with TEV protease, and dialyzed into buffer containing 25 mM Na_2_HPO_4_ [pH 8.0], 400 mM NaCl, 0.4 mM BME at 4ºC overnight. The next day, dialysate was recirculated over a 5 mL HiTrap chelating column to capture the his6-MBP-TEV tag. The flow through containing NF1(1-333aa) was Vivaspin 20 (5 kDa MWCO) and loaded onto a Superdex 75 column equilibrated in buffer 20 mM HEPES [pH 7], 200 mM NaCl, 10% glycerol, 1 mM TCEP. Peak fractions containing monodispersed NF1 were pooled, concentrated to 250 µM, and snap frozen in liquid nitrogen prior to storage at -80ºC.

#### GEF(fb)

A gene sequence encoding GEF(fb, feedback) chimeric protein was expressed into BL21 *E. coli* as a his_10_-TEV-SUMO3-SosCat(564-1049aa)-SNAP-RBD fusion protein. Bacteria were grown at 37°C in Terrific Broth to OD_600_ = 0.8. Culture was shifted to 27°C for 1 hour then induced with 0.1 mM IPTG. Expression was allowed to continue for 13 hours before harvesting. Cells were lysed in 50 mM Na_2_HPO_4_ [pH 8.0], 400 mM NaCl, 0.4 mM BME, 1 mM PMSF, 100 μg/mL DNase by sonication. Lysate was centrifuged at 15,000 rpm (35000 x g) for 60 minutes in a Beckman JA-20 rotor at 4ºC. Lysate was circulated over a 5 mL HiTrap chelating column (Cytiva GE17-0409-01) inoculated with 100 mM CoCl_2_ for ten minutes. Column was washed with 50 mM Na_2_HPO_4_ [pH 8.0], 400 mM NaCl, 10 mM imidazole, 0.4 mM BME. Bound protein was eluted in 50 mM Na_2_HPO_4_ [pH 8.0], 400 mM NaCl, 500 mM imidazole, 0.4 mM BME. Peak fractions were pooled and a his-tagged SenP2 SUMO protease was added to cleave off the SUMO3 tag. The solution was dialyzed against 4 liters of buffer containing 20 mM HEPES [pH 7.5], 400 mM NaCl, 5% glycerol, 0.4 mM BME. Dialysate containing cleaved protein was recirculated over the HiTrap chelating column for 90 minutes. Flowthrough was concentrated in a 30 kDa MWCO Amicon-Ultra concentrator before loading onto a desalting column equilibrated in 20mM Tris [pH 8.0], 150 mM NaCl, 10% glycerol, 1 mM TCEP. Peak fractions were pooled and snap frozen in liquid nitrogen for storage at -80ºC.

#### GEF(ff)

The protein referred to as GEF (ff, feed-forward) represents a PIP_3_ lipid binding domain derived for Btk fused to the catalytic domain of Sos1(564-1049aa). The resulting chimeric protein was expressed as a gene sequence encoding his_6_-SUMO-Btk(R49S/K52S)-Sos1(564-1049aa) in BL21(DE3) *E. coli* that contain vectors for constitutive expression for Trigger factor and rare tRNAs (i.e. pRIL). Bacteria were grown at 37°C in Terrific Broth until it reached OD_600_ = 0.8. Cultures were shifted to 18°C for 1 hour then induced with 0.1 mM IPTG. Expression was allowed to continue for 18 hours before harvesting. Cells were lysed in 50 mM Na_2_HPO_4_ [pH 8.0], 400 mM NaCl, 10 mM imidazole, 4 mM BME, 1 mM PMSF, 100 μg/mL DNase by sonication. Lysate was centrifuged at 15,000 rpm (35000 x *g*) for 60 minutes in a Beckman JA-20 rotor at 4ºC. NiNTA resin was added to the supernatant and allowed to bind for 1 hour, then washed with 50 mM Na_2_HPO_4_ [pH 8.0], 400 mM NaCl, 10 mM imidazole, 4 mM BME. Bound protein was eluted in 50 mM Na_2_HPO_4_ [pH 8.0], 400 mM NaCl, 500 mM imidazole, 4 mM BME. Peak fractions were pooled and an Ulp1 protease was added to cleave off the his_6_-SUMO tag. The solution was dialyzed against 4 liters of buffer containing 50 mM Na_2_HPO_4_ [pH 8.0], 300 mM NaCl, 20 mM imidazole, 5% glycerol, 0.4 mM BME. Dialysate containing cleaved protein was recirculated over NiNTA resin. Flowthrough was loaded onto a G25 Sephadex desalting column equilibrated in 20 mM HEPES [pH 7.0], 50 mM NaCl, 1 mM TCEP. Peak fractions were pooled and loaded onto a HiTrap Capto S cation exchange column. The Btk1-GEF eluted in the presence of 450 mM NaCl. These fractions were concentrated in a 50 kDa MWCO Amicon-Ultra concentrator and snap frozen in liquid nitrogen for storage at -80ºC.

#### p110γ(PIK3CG), ybbr-p101(PIK3R5)

High five insect cells were infected with baculovirus to drive the dual expression of human p110γ(PIK3CG, 1-1102aa) and TwinStrept-his10-TEV-ybbr-p101(PIK3R5, 1-880aa) from polyhedrin promoters are previously described ^15^. The ybbR13 motif was following peptide sequence: D<u>S</u>LEFIASKLA ^88^. BACMIDs and baculovirus were generated following protocols previously described ^89^. Optimal protein expression conditions were empirically determined to minimize cell death and maximize protein yield (typically 1.5-2% vol/vol final concentration of P2 virus). After 48 hrs of protein expression, cells were harvested and pellets were frozen at -80°C. The PI3Kγ complex was purified using previously published protocols ^90^. In brief, cells were lysed in 20 mM Tris [pH 8.0], 100 mM NaCl, 10 mM imidazole, 5% glycerol, 2 mM BME, 1 mM PMSF, 100 μg/mL DNase by sonication. After lysis, Triton-X was added to a final concentration of 0.1%. Lysate was centrifuged at 30,000 rpm for 60 minutes in a Beckman TI-45 rotor at 4ºC. NiNTA resin was added to the supernatant and allowed to bind for 1 hour, then washed with 20 mM Tris [pH 8.0], 100 mM NaCl, 10 mM imidazole, 5% glycerol, 2 mM BME. Bound protein was eluted in 20 mM Tris [pH 8.0], 100 mM NaCl, 500 mM imidazole, 5% glycerol, 2 mM BME. Peak fractions were pooled, bound to Strep-Tactin Sepharose resin (IBA 2-1201-002), and washed with 100 mM Tris [pH 8.0], 150 mM NaCl, 1 mM EDTA. Bound protein was eluted in 100 mM Tris [pH 8.0], 150 mM NaCl, 1 mM EDTA, 2.5 mM desthiobiotin (Sigma, Cat #D1411) and a his_6_-tagged TEV protease was added to cleave the his6 affinity tag off the catalytic subunit. The solution was dialyzed against 4 liters of buffer containing 20 mM Tris [pH 8.0], 150 mM NaCl, 10% glycerol, 1 mM TCEP. The dialysate was concentrated in a 100 kDa MWCO Amicon-Ultra concentrator and loaded on a Superdex 200 gel filtration column equilibrated in 20 mM Tris [pH 8.0], 150 mM NaCl, 10% glycerol, 10% glycerol, 1 mM TCEP. Peak fractions were pooled and snap frozen in liquid nitrogen for storage at -80ºC. To fluorescent label the PI3Kγ complex (p110γ/ybbr-p101), frozen protein was thawed and combined with Sfp transferase and Dyomics647-CoA (Dy647-CoA) as previously described ^90^. The final labeled PI3Kγ complex was separated from unreacted DY647-CoA and Sfp using Superdex 200 gel filtration column equilibrated in 20 mM Tris [pH 8], 150 mM NaCl, 10% glycerol, 1 mM TCEP, 0.05% CHAPS. Peak fractions were pooled and concentrated to 5-10 µM before flash freezing with liquid nitrogen and stored in -80ºC freezer.

#### Btk-SNAP

The PI(3,4,5)P_3_ binding domain derived from Btk was recombinantly expressed and purified as a his6-SUMO-Btk(PH-TH,R49S/K52S)-SNAP fusion protein using a protocol previously described ^91^. Purified SNAP-RBD and Btk(PH-TH,R49S/K52S)-SNAP were labeled in vitro by combining 1.5x molar excess of SNAP-Surface Alexa546 dye (NEB, Cat# S9132S) or SNAP-Surface Alexa647 dye (NEB, Cat# S9136S) with purified proteins. Labeling reactions were incubated overnight at 4ºC in buffer containing 20 mM Tris [pH 8.0], 200 mM NaCl, 1 mM TCEP. The next day, labeled proteins were desalted into buffer containing 20 mM Tris [pH 8.0], 200 mM NaCl, 10% glycerol, 1 mM TCEP using a PD10 column to remove most free SNAP dye. Proteins were concentrated in an Amicon filter and loaded onto a Superdex 75 column equilibrated with 20 mM Tris [pH 8.0], 200 mM NaCl, 10% glycerol, 1 mM TCEP. Unreacted SNAP dyes and labeled proteins were separated by SEC. The peak elution was pooled, concentrated, aliquoted, and flash frozen with liquid nitrogen.

#### Farnesylated GβGγ Complex

Sf9 insect cells were infected with baculovirus (2% vol/vol) to drive the dual expression of his6-TEV-Gγ_2_ (1-71aa) and Gβ_1_ (1-340aa) polyhedrin promoters as previously described ^91^. After 48 hours of expression at 27°C, cells were harvested and pellets were frozen at -80°C. Cells were lysed in 50 mM HEPES [pH 8.0], 100 mM NaCl, 3 mM MgCl_2_, 0.1 mM EDTA [pH 8.0], 10 µM GDP, 10 mM BME, Sigma PI tablets, 1 mM PMSF, 100 μg/mL DNase by sonication and centrifuged at 800 x *g* at 4ºC. The supernatant was centrifuged at 100,000 x *g* for 30 minutes in a Beckman TI-70 rotor at 4ºC. The pellet was resuspended in 50 mM HEPES [pH 8.0], 50 mM NaCl, 3 mM MgCl_2_, 1% sodium cholate (wt/vol), 10 µM GDP, 10 mM BME, 1 mM PMSF, dounce homogenized, stirred for 1 hour, and centrifuged at 100,000 x *g* for 45 minutes in a Beckman TI-70 rotor at 4ºC. The supernatant was diluted 5-fold in 20 mM HEPES [pH 7.7], 100 mM NaCl, 0.1 % C_12_E_10_ (Polyoxyethylene (10) lauryl ether), 20 mM imidazole, 2 mM BME and allowed to bind to NiNTA resin for 2 hours. The resin was washed with 20 mM HEPES [pH 7.7], 100 mM NaCl, 0.1 % C_12_E_10_, 20 mM imidazole, 2 mM BME. The Gα subunit was eluted in 20 mM HEPES [pH 7.7], 100 mM NaCl, 0.1 % C_12_E_10_, 20 mM imidazole, 2 mM BME, 50 mM MgCl_2_, 10µM GDP, 30 µM AlCl_3_, 10 mM NaF at 30ºC. The complex was eluted in 20 mM Tris [pH 8.0], 25 mM NaCl, 0.1 % C_12_E_10_, 200 mM imidazole, 2 mM BME and a his_6_TEV protease was added to cleave the his_6_ tag off the Gγ_2_ subunit. The solution was loaded onto a G25 Sephadex desalting column equilibrated in 20 mM Tris-HCl [pH 8.0], 25 mM NaCl, 8 mM CHAPS, 2 mM TCEP. Peak fractions were pooled and loaded onto a MonoQ anion exchange column. The GβGγ complex eluted at ∼180 mM NaCl. These fractions were concentrated in a 30 kDa MWCO Amicon-Ultra concentrator and loaded on a Superdex 75 gel filtration column equilibrated in 20 mM Tris [pH 8.0], 100 mM NaCl, 8 mM CHAPS, 2 mM TCEP. Peak fractions were pooled and snap frozen in liquid nitrogen for storage at -80ºC.

#### PTEN

PTEN was expressed in High 5 insect cells as a PTEN(1-352aa)-GGLPETGGGY-MxeGyrA-his6. Although not utilized in this study, the LPETGG motif was added for sortase mediated peptide ligation. The tyrosine positioned N-terminal to the MxeGyrA intein increases the efficiency of DTT induced intein cleavage. High cells expressing PTEN were lysed by Dounce homogenization in buffer containing 50 mM Na_2_HPO_4_ [pH 8.0], 10 mM imidazole, 300 mM NaCl, 0.1% Triton X-100, 0.5 mM ABESF, DNase, Roche protease inhibitor tablet (1 per 100 mL of lysis buffer), and no reducing agent. Lysate was clarified by centrifugation 35,000 rpm in Beckman Ti-70 rotor for 45 minutes. Clarified cell lysate was incubate with 5 mL of NiNTA resin in a beaker with stir bar at 4ºC for 1 hour. Resin was washed with 100 mL of buffer containing 50 mM Na_2_HPO_4_ [pH 8.0], 10 mM imidazole, 400 mM NaCl. PTEN(1-352aa)-GGLPETGGGY-MxeGyrA-his6 was eluted from the NiNTA resin with 20 mL of 50 mM Na_2_HPO_4_ [pH 8.0], 500 mM imidazole, 400 mM NaCl. The MxeGyrA-his6 was cleaved off by incubating with 10 mM DTT overnight at 4ºC. To separate PTEN(1-352aa)-GGLPETGGGY (pI = 8.18) and MxeGyrA-his6 (pI = 5.81), the cleaved protein was desalted into buffer containing 20 mM HEPES [pH 7.0], 50 mM NaCl, 2 mM DTT. The proteins were then load on a MonoS column equilibrated in 20 mM HEPES [pH 7.0], 50 mM NaCl, 2 mM DTT and resolved with a 5-100% salt gradient (100% = 20 mM HEPES [pH 7.0], 1 M NaCl). PTEN eluted from the MonoS column in the presence of 500 mM NaCl at pH 7.0. Peak fractions were pooled and concentrated with VivaSpin 6 (10 kDa MWCO). PTEN was loaded on a Superdex 75 in buffer containing 20 mM Tris [pH 8.5], 200 mM NaCl, 10% glycerol, 0.5 mM TCEP. Peak fractions were pooled, concentrated to 112 µM, and flash frozen with liquid nitrogen before being stored in -80ºC.

#### Preparation of supported lipid bilayers

The following lipids were used to generate small unilamellar vesicles (SUVs) and subsequently supported lipid bilayers: 1,2-dioleoyl-sn-glycero-3-phosphocholine (18:1 DOPC, Avanti #850375C), L-α-phosphatidylinositol-4,5-bisphosphate (Brain PIP_2_, Avanti #840046X), 1,2-dioleoyl-sn-glycero-3-phosphoethanolamine-N-[4-(p-maleimidomethyl) cyclohexane-carboxamide] (18:1 MCC-PE, Avanti #780201C), 1,2-dioleoyl-sn-glycero-3-[(N-(5-amino-1-carboxypentyl)iminodiacetic acid)succinyl] (18:1 DGS-NTA(Ni), Avant #790404C), egg L-α-Phosphatidylethanolamine-N-(lissamine rhodamine B sulfonyl) (Ammonium Salt) (Liss Rhod PE, Avanti # 810146). Throughout the paper PI(4,5)P_2_ and PI(3,4,5)P_3_ are referred to as PIP_2_ and PIP_3_, respectively. To make SUVs, 2 µmoles total lipids were combined in a 35 mL glass round bottom flask containing ∼2 mL of chloroform.

Lipids were dried to a thin film using rotary evaporation with the glass round-bottom flask submerged in a 42ºC water bath. After evaporating the chloroform, the round bottom flask was flushed with nitrogen gas or placed in a vacuum desiccator. The lipid film was resuspended in 2 mL of PBS [pH 7.4] to make a final concentration of 1 mM total lipids. All lipid mixtures expressed as percentages (e.g. 98% DOPC, 2% PIP_2_) are equivalent to molar fractions. To generate 50 nm SUVs, 1 mM total lipid mixtures were extruded through a 0.05 µm pore size 19 mm polycarbonate membrane (Whatman, Cat #WHA800308) with filter supports (Avanti #610014) on both sides of the PC membrane. To create supported lipid bilayers, coverglass (25x75 mm, IBIDI, Cat #10812) was initially cleaned with 2% Hellmanex III (Fisher, Cat #14-385-864) that was heated to 60-70ºC in a glass coplin jar and incubated for at least 30 minutes. After rinsing the Hellmanex cleaned glass with MilliQ water, the coverglass was etched with Piranha solution (1:3, hydrogen peroxide:sulfuric acid) for 10-15 minutes. Etched coverglass that was rinsed and stored in MilliQ water, were individually removed from the Coplin jar and rapidly dried with nitrogen gas before adhering to a 6-well, sticky-side chamber (IBIDI, Cat #80608). To create a supported lipid bilayer, a total lipid concentration of 0.25 mM SUVs in 1x PBS [pH 7.4] was flowed into the IBIDI chamber. After 30 minutes of incubation, supported membranes were washed with 5 mL of PBS [pH 7.4] to remove non-absorbed SUVs. Membrane defects were then blocked for ∼5 minutes with 1 mg/mL beta casein (ThermoFisher, Cat #37528) diluted in 1x PBS [pH 7.4]. To reduce the fraction of immobile proteins on SLBs, the 10 mg/mL beta casein solution was centrifuged in the TLA120.2 at 100,000 rpm (434,513 x *g*) for 30 minutes at 4ºC using an Optima MAX-TL ultracentrifuge. After blocking bilayers with beta casein, membranes were washed again with 2 mL of PBS and then stored at room temperature for up to 2 hours before mounting on the microscope. Prior to single molecule imaging experiments, supported membranes were washed into TIRF imaging buffer.

#### Membrane conjugation

For experiments involving membrane conjugated H-Ras, beta casein blocked SLBs containing MCC-PE lipids were incubated for 2 hours with 30-35 µM H-Ras at 23ºC in buffer containing 1x PBS [pH 7.4], 1 mM MgCl_2_, 100 µM GDP, 0.1 mM TCEP. Bilayers were quenched in 1x PBS [pH 7.4], 5 mM 2-mercaptoethanol for 10 minutes before washing with 1x PBS [pH 7.4], 1 mM MgCl_2_, 100 µM GDP, 0.1 mM TCEP. The resulting Ras membrane density ranged from 1000-2000 molecules/µm^2^ as previously described ^32^. Following conjugation of H-Ras, membranes were incubated with 20-50 nM his_10_ -mNG-iLID for 20 minutes at 23ºC. Unbound his_10_-mNG-iLID was flushed out of the chamber with TIRF-M imaging buffer. For simplicity, his_10_-mNG-iLID is referred to as iLID throughout the manuscript. For experiments involving GβGγ (i.e. Figure 6), bilayers were incubated with 25 nM GβGγ (native farnesyl lipid anchor) for 20 minutes. Unbound GβGγ was flushed out of the chamber with TIRF-M imaging buffer.

#### TIRF microscopy imaging buffer

All supported lipid bilayer TIRF-M experiments were performed using the following reaction buffer: 20 mM HEPES [pH 7.0], 150 mM NaCl, 1 mM ATP (Sigma, Cat #A2383-10G), 100 µM GTP, 5 mM MgCl_2_, 0.5 mM EGTA, 20 mM glucose, 200 µg/mL beta casein (ThermoScientific, Cat #37528), 20 mM BME, 320 µg/mL glucose oxidase (Serva, Cat #22780.01 *Aspergillus niger*), 50 µg/mL catalase (Sigma, Cat #C40-100MG Bovine Liver), and 2 mM Trolox (Cayman Chemicals, Cat #10011659). For experiments shown in **Fig. 6b** and **Supplemental Fig. 7b**, ATP was initially omitted from buffer and 1 mM ATP was spiked in at indicated timepoint. Perishable reagents (i.e. glucose oxidase, catalase, and Trolox) were added 5-10 minutes before initiating biochemical reconstitutions on SLBs and acquiring images by TIRF-M. All reactions in this study were reconstituted and visualized on a TIRF microscope at 23ºC.

#### Microscope hardware and imaging acquisition

Imaging experiments were performed using an inverted Nikon Ti2 microscope using either a 60x (1.49 NA) or 100x (1.49 NA) oil immersion Nikon TIRF objective. The x-axis and y-axis positions were controlled using a Nikon motorized stage, joystick, and NIS element software. Fluorescently labeled proteins were excited with either a 488, 561, or 637 nm OBIS laser diode (Coherent Inc. Santa Clara, CA) controlled with a Vortran laser launch (Sacramento, CA) and acousto-optic tuneable filters (AOTF) control. Excitation and emission light was transmitted through a multi-bandpass quad filter cube (C-TIRF ULTRA HI S/N QUAD 405/488/561/638; Semrock) containing a dichroic mirror. For bulk excitation of fluorescent biosensors, laser power measured through a 60x TIRF objective ranged from 0.5-2 mW. Fluorescence emission was detected on an iXion Life 897 EMCCD camera (Andor Technology Ltd., UK) after passing through a Nikon Ti2 emission filter wheel containing the following 25 mm emission filters: ET525/50M, ET600/50M, ET700/75M (Semrock). All experiments were performed at room temperature (23ºC). Microscope hardware was controlled with Nikon NIS elements.

#### Spatially patterned light

A Mightex Polygon400 digital micromirror device (DMD) controlled by NIS elements was used for patterned illumination. For activation of membrane bound iLID, excitation light from a SOLA SE light engine (Lumencor, 380-680 nm range of wavelengths) was filtered using a Chroma 488 nm long-pass dichroic mirror (25.5 x 36 x 2 mm; Cat #ZT488rdc-UF2). The dichroic mirror was mounted at 45º angle in a Nikon Ti2 LAPP (Cat #Ti2-LA-BM-E) prior to entering the microscope filter turret. This setup reflects iLID activating light from the Polygon400 DMD to the sample. The 488LP dichroic allowed transmission of 561 nm and 647 nm laser excitation light to image fluorescently labeled SspB, GEFs, and activity biosensors following activation of iLID. Following repetitive illumination of the same membrane region with 488 nm light, we observed a very slight increase in light-independent basal SspB recruitment. This potentially represents an accumulation of iLID protein that does not transition back to its “closed”, dark-state conformation. For this reason, we changed the field of view between light-induced activation steps in the same reaction chamber for subsequent experiments.

### Fisher wave simulation

A Fisher wave is a propagating wave that arises in reaction-diffusion systems that combine diffusion and activation. The specific equation we used to model the amplification and expansion of Ras(GTP) in the presence of global inhibitors follows:

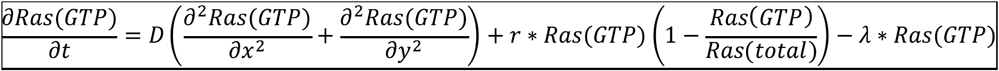

The chemical concentration of Ras(GTP) at position x and time t was defined as 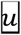 (x,t) or Ras(GTP)(x,t). Variables in the above equation are defined as follows: D is the diffusion coefficient of Ras(GTP) in µm^2^/second, r is the intrinsic growth rate (i.e. catalysis), and λ is the decay rate (i.e. RasGAP activity). The maximum or total Ras density (*K*) was 20-100 molecules/µm^2^. The term 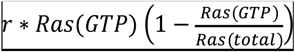 represents a logistic growth model in which the growth rate decreases as Ras(GTP) approaches Ras(total). Refer to Supplementary Table 1 for specific parameters used to generate data in the main figures. Python code required to run the Fisher wave simulations is appended as a Supplementary .py file. Python code was executed using Spyder 6 software.

#### Image analysis

To calculate the fold membrane enrichment of SspB fusion proteins (**Fig. 2h, i** and **Supplemental Fig. 4g**) we took into account both the basal level of light-independent recruitment and the maximum light-induced membrane recruitment, as previously described ^92^. Fold membrane enrichment of AF647-SspB fusion proteins was calculated using (basal light-independent recruitment + peak 488 nm light dependent recruitment)/ basal light-independent recruitment. The same equation was used to calculate fold Ras activation in **Fig. 3e, f**: (steady-state light-independent Ras activation + peak 488 nm light dependent activation)/steady-state light-independent activation

To calculate the decay halftime of AF647-SspB membrane dissociation (**Fig. 2h, j** and **Supplemental Fig. 4h**), kinetic traces were fit to a non-linear one phase decay curve using Prism GraphPad 10: Y = (Y0 - Plateau)*exp(-K*X) + Plateau where Y0 is the Y value at time 0, Plateau is the Y value at infinite times, and K is the rate constant, expressed in reciprocal of the X axis time units.

Ras(GTP) wave radius (µm) (**Fig. 4f**) was calculated as a function of time based on AF546-RBD fluorescence using the manual thresholding Fiji/ImageJ plugin ^93^. Starting with an 8-bit image stack, images were globally thresholded at a pixel value of 40. Outlier fluorescent speckles caused by immobilized proteins and membrane defects smaller than 9 pixels were removed. The images were subsequently eroded using Fiji/ImageJ. These thresholded image stacks were further analyzed using Python code executed in Spyder 6 software. For each image in the time stack, radii were measured from the centroid of the Ras(GTP) region at 15 degree intervals (24 total) and averaged.

To generate the intensity line scans in **Fig. 5**, images were first blurred in Fiji/ImageJ ^93^ using a Gaussian equation with a variance of 1.5 µm. Intensities were measured along a horizontal line at the center of each image. To determine the steepness of Fisher waves (**Fig. 6h**) we calculated the derivative at the wave front inflection point using Prism GraphPad 10.

