## Supplemental Figures for "Reconstitution of Ras-PI3Kγ membrane communication and feedback using light-induced signaling inputs"

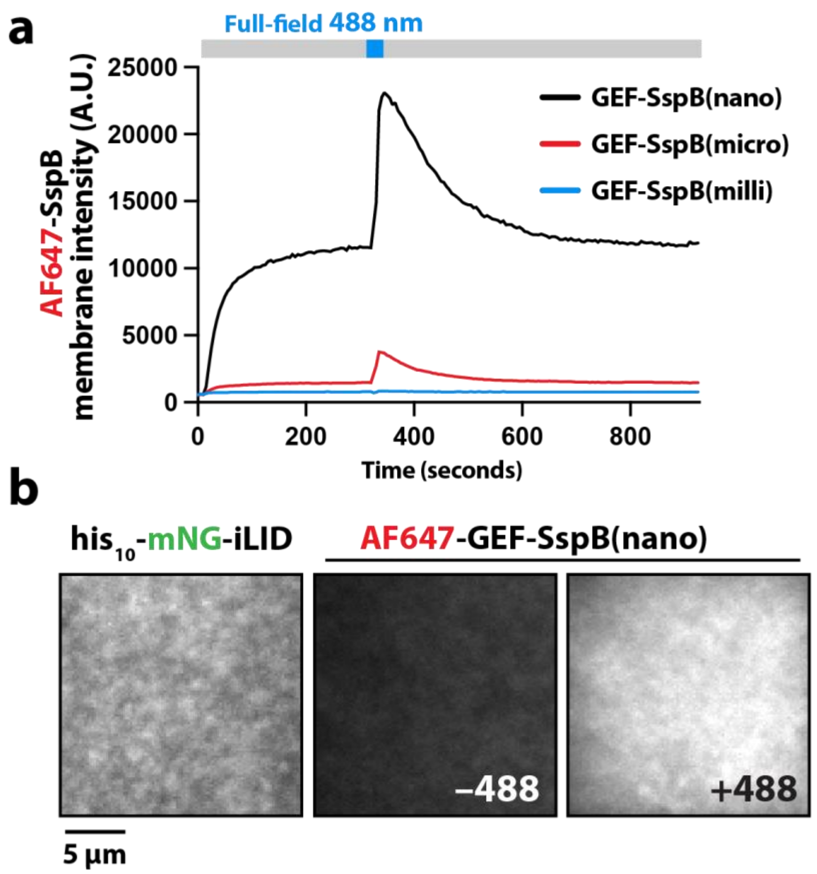


**Supplementary Figure 1**

**Light-induced heterodimerization of iLID and GEF-SspB on supported lipid bilayers**

**(a)** AF647-GEF-SspB variants (nano, micro, and milli) have distinct membrane absorption kinetics in the absence of 488 nm light and exhibit different magnitudes of membrane recruitment following a uniform, 50 ms pulse of 488 nm laser light. All variants exponentially dissociate from the membrane in the absence of 488 nm light. Membranes were incubated with 50 nM $his_{10}$-mNG-iLID for 20 minutes. Unbound $his_{10}$-mNG-iLID was washed out of sample chamber prior to flowing in 50 nM AF647-GEF-SspB. **(b)** Representative TIRF-M images showing the membrane localization of membrane-tethered $his_{10}$-mNG-iLID and AF647-GEF-SspB(nano) in the absence and presence of 488 nm light. Membrane composition: 96% DOPC, 2% NiNTA (+iLID bound), 2% MCC-PE lipids (+Ras bound).

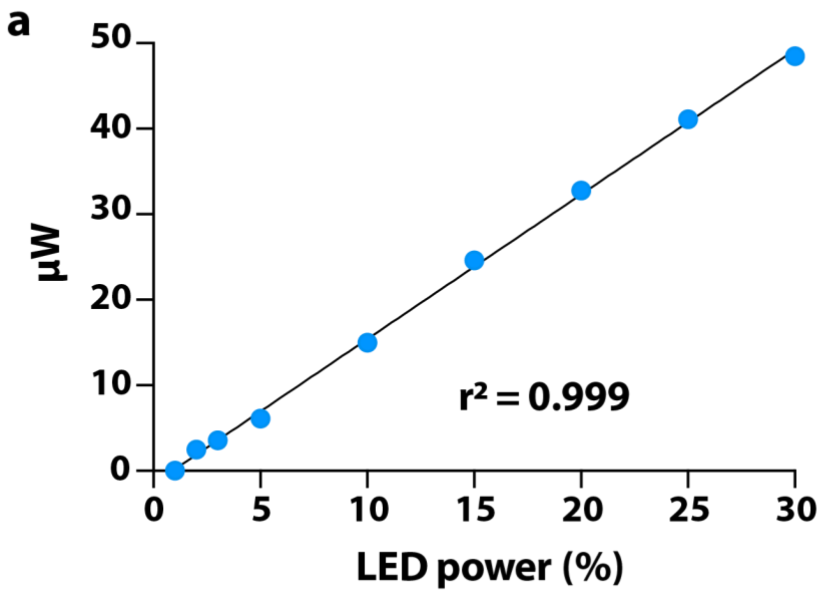


**Supplementary Figure 2**
**Calibration of LED power for iLID activation**
**(a)** Intensity of 488 nm light (μW) measured through a 60x TIRF objective as a function of the LED power (%) from SOLA SE light engine. Intensity was measured using a Newport power meter positioned in place of the glass coverslip.

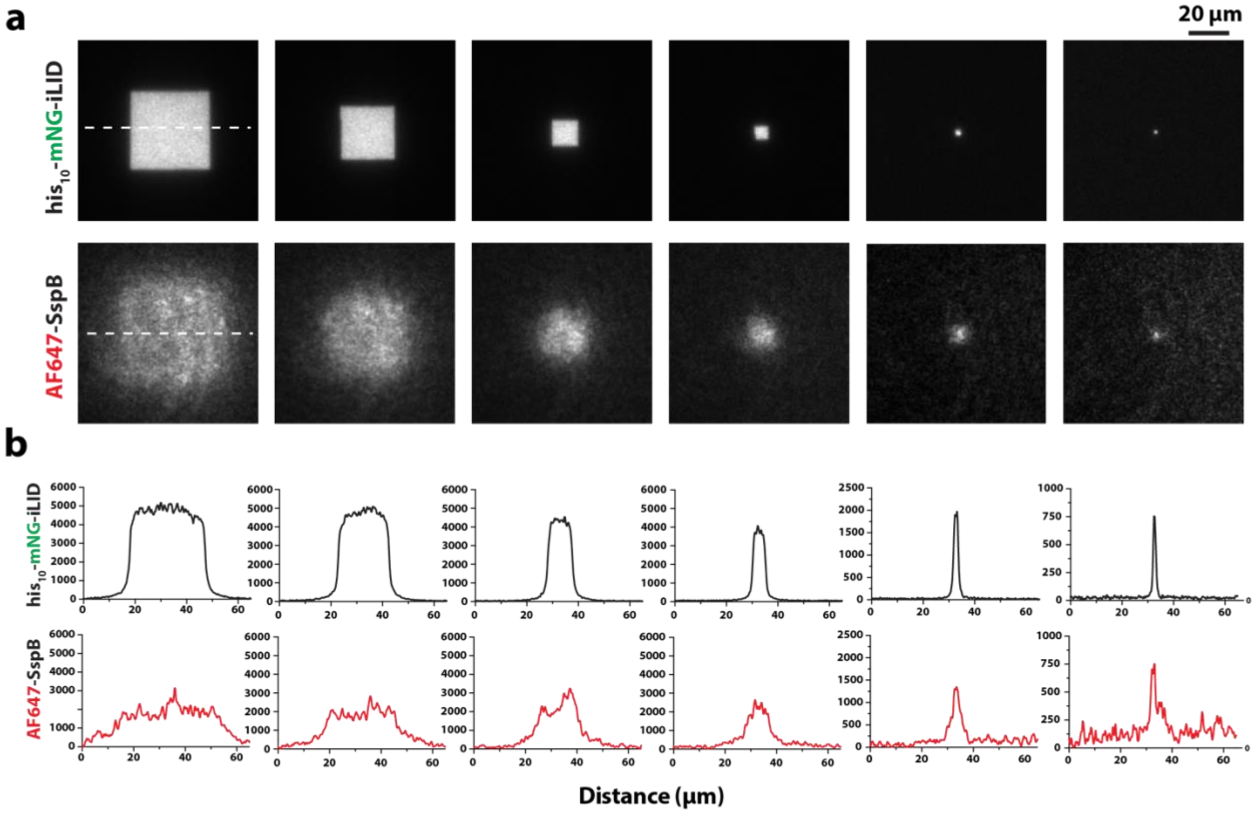


**Supplementary Figure 3**

**SspB membrane recruitment as a function of pattern size**

**(a)** TIRF-M images showing 488 nm-excited his$_{10}$-mNG-iLID in squares of different side lengths (900, 400, 100, 25, 4, 1 µm$^2$) on bilayers and corresponding AF647-SspB(micro) membrane localization. **(b)** Line scans showing fluorescence intensity of membrane-tethered his$_{10}$-mNG-iLID and membrane-recruited AF647-SspB(micro) 5 seconds after a 488 nm pulse of light. A representative line scan (dashed line) is shown in **(a)**. **(a, b)** Membrane composition: 96% DOPC, 2% NiNTA (+iLID bound), 2% MCC-PE lipids. All experiments contained a solution concentration of 100 nM AF647-SspB(micro).

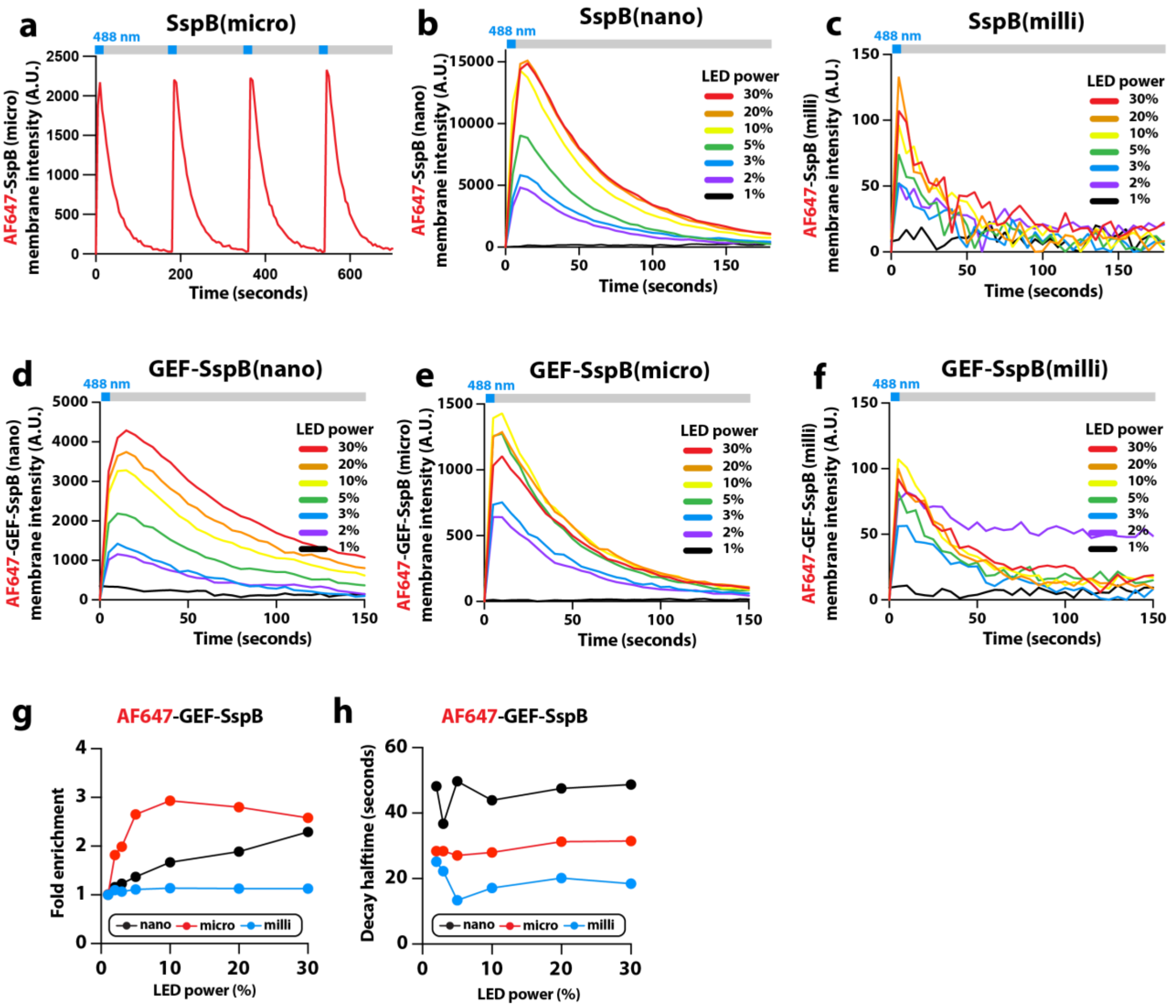


**Supplementary Figure 4**

**Kinetics of membrane recruitment and dissociation for SspB variants**

**(a)** Repeated light-dependent membrane recruitment of 50 nM AF647-SspB(micro) to the same 20 μm-diameter circle. **(b, c)** Kinetic traces showing membrane recruitment and dissociation of 50 nM AF647-SspB (nano and milli) measured using a 488 nm light pulse at a range of intensities (1-30% LED power, 0-50 μW, conversion in **Supplementary Fig. 2a**). **(d-f)** Membrane recruitment and dissociation kinetics using a 488 nm light pulse at a range of intensities (1-30% LED power, 0-50 μW, conversion in **Supplementary Fig. 2a**) in the presence of **(d)** 50 nM AF647-GEF-SspB(nano), **(e)** 50 nM AF647-GEF-SspB(micro), and **(f)** 50 nM AF647-GEF-SspB(milli). **(g)** Quantification of AF647-GEF-SspB(nano, micro, and milli) fold membrane enrichment as a function of light power. Fold enrichment values were calculated using (basal recruitment + 488 nm light recruitment)/basal recruitment as described in **Fig. 2h. (h)** Quantification of AF647-GEF-SspB(nano, micro, and milli) signal decay half-time ($t_{1/2}$) as a function of light power. $t_{1/2}$ values were calculated as described in **Fig. 2h.**

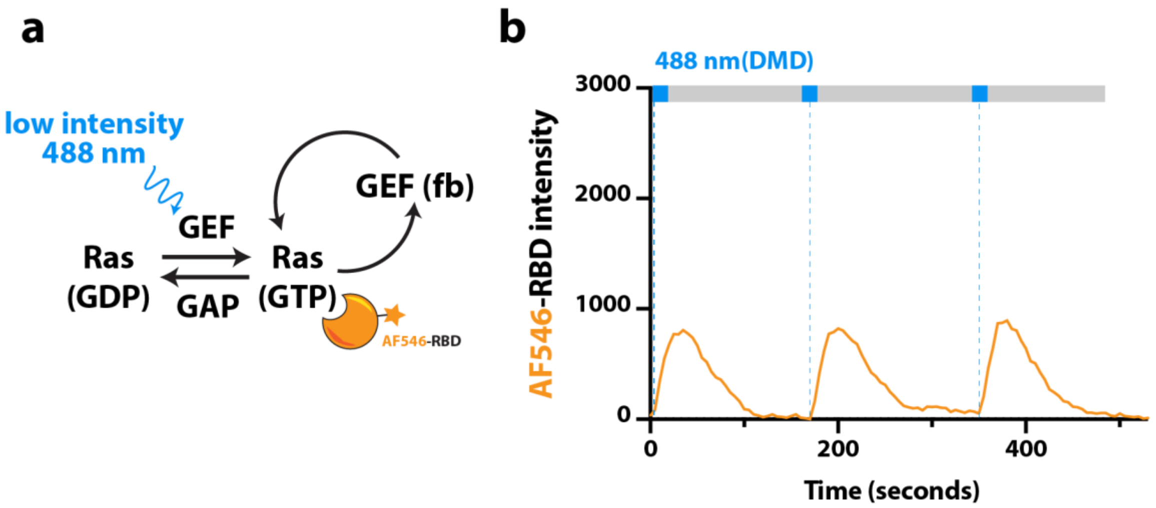


**Supplementary Figure 5**

**Repetitive low light induced membrane recruitment of GEF-SspB does not trigger Ras(GTP) positive feedback**

**(a)** Time delayed membrane recruitment of AF647-GEF-SspB(micro) does not trigger Ras(GTP) positive feedback when fold enrichment is too low. Reactions contained 10 nM GEF(fb), 10 nM AF647-GEF-SspB(micro), 544 nM RasGAP, and 50 nM AF546-RBD. Membrane composition: 92% DOPC, 2% NiNTA (+iLID bound), 2% MCC-PE (+Ras bound), and 4% $PIP_2$.

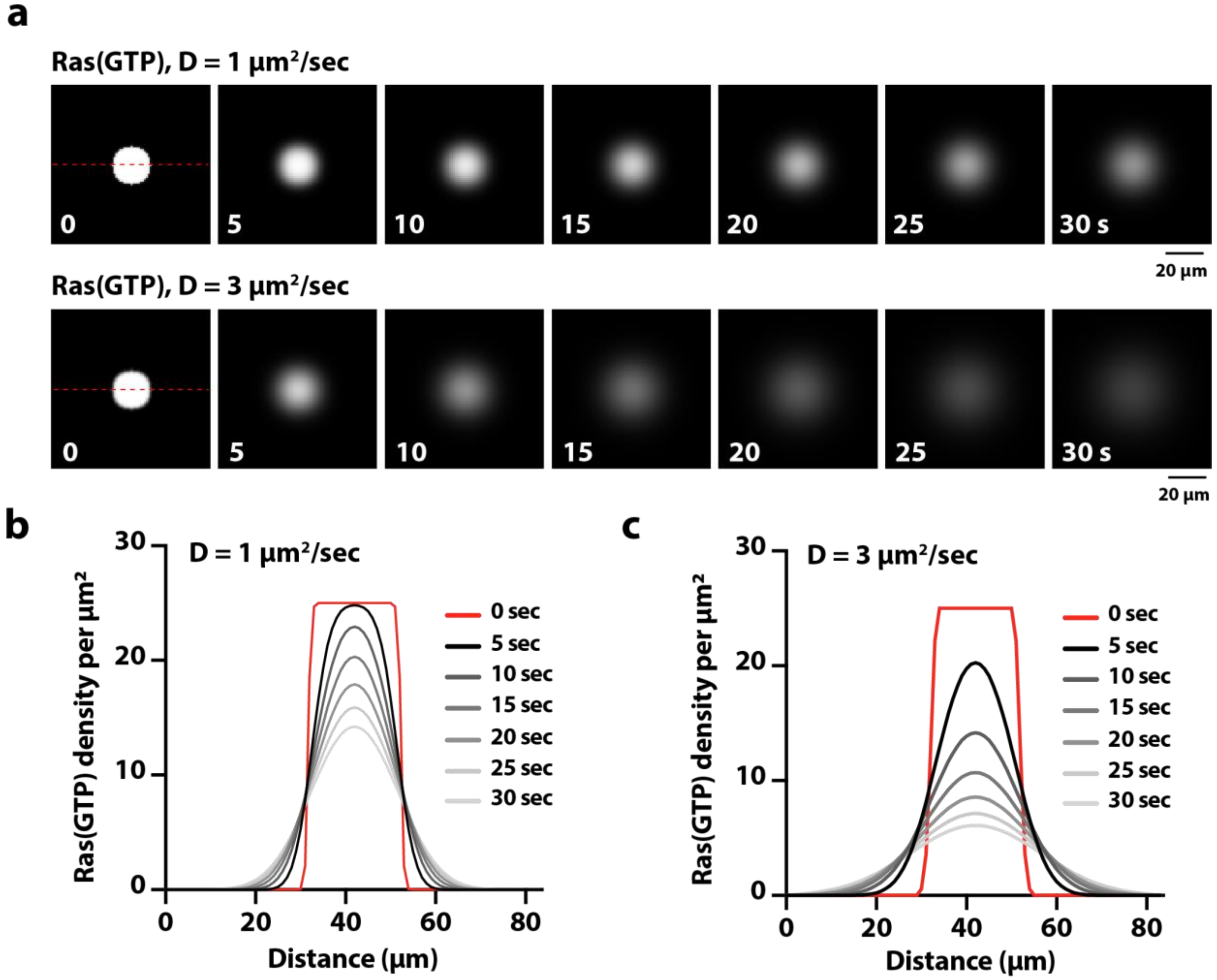


**Supplementary Figure 6**

**Fisher waves simulated in the absence of logistical growth and GAP-mediated decay**

**(a)** Montages showing the change in Ras(GTP) membrane intensity for Fisher wave simulations lacking positive feedback and GAP-mediated decay. Reactions were simulated with Ras(GTP) having diffusion coefficients of 1 or 3 $\mu m^2$/sec. Red dashed lines represent the position of line scans plotted below. **(b-c)** The distance Ras(GTP) diffuses from original site of activation as a function of time, depends on the diffusion coefficient. Line scans showing the Ras(GTP) membrane density (molecules/$\mu m^2$) as a function of the membrane position for simulations performed with **(b)** 1 $\mu m^2$/sec or **(c)** 3 $\mu m^2$/sec. Legends indicate time following initial Ras activation (solid red line) which seeds 25 Ras(GTP)/$\mu m^2$.

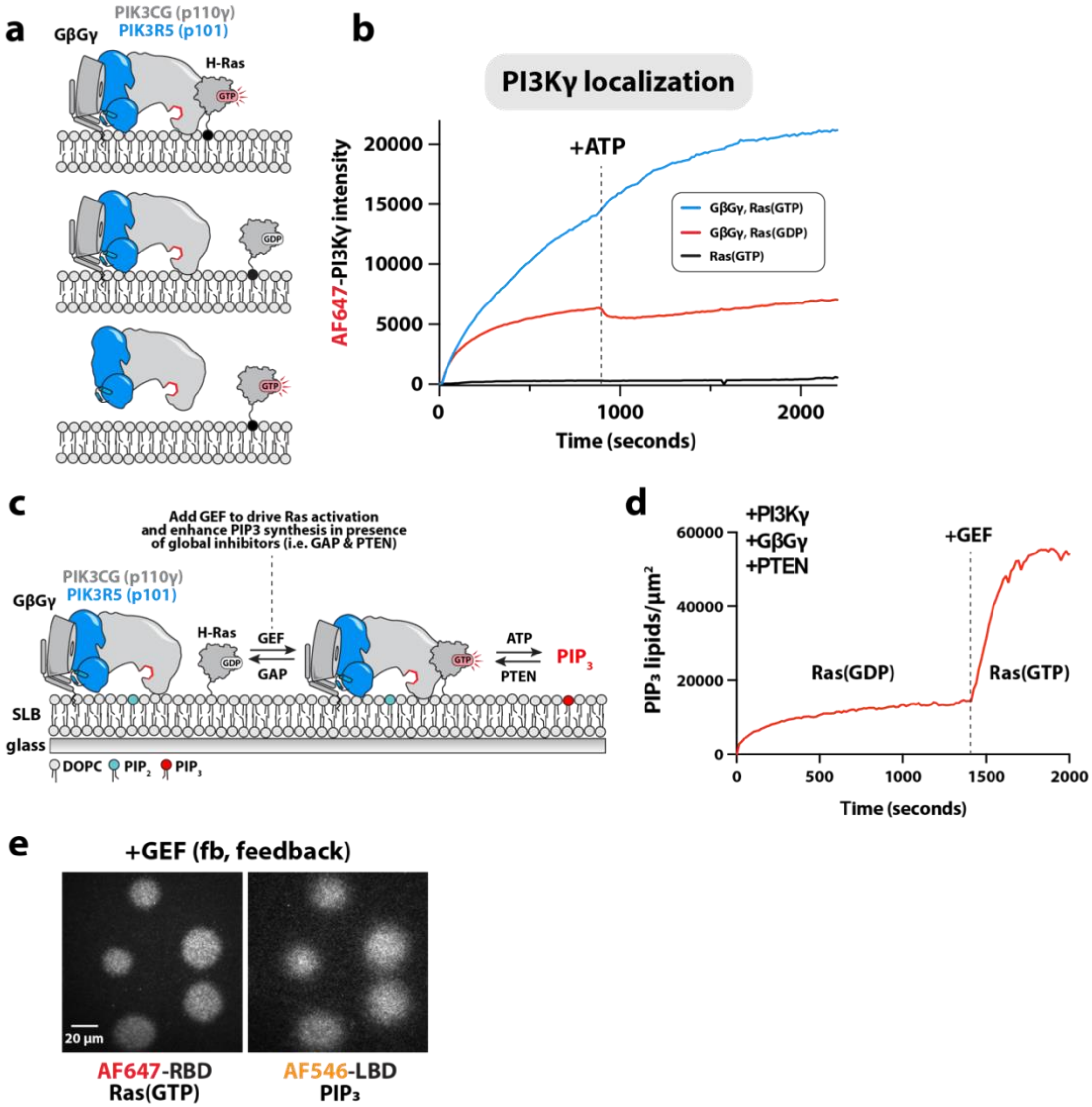


**Supplementary Figure 7**

**Reconstitution of Ras(GTP)-PI3Kγ communication and feedback**

**(a)** Experimental design for reconstitution of Ras(GTP)-PI3Kγ communication and feedback on supported lipid bilayer. **(b)** Membrane localization kinetics of AF647-PI3Kγ in the presence of Ras(GTP), GβGγ, or both inputs. Dashed line marks the addition of 1 mM ATP. **(c)** The presence of both GβGγ and Ras(GTP) allows PI3Kγ activity to overcome PTEN lipid phosphatase activity. $PIP_3$ production increases rapidly when Ras is activated by the addition of 50 nM GEF (dashed line). **(b-c)** Membrane composition: 94% DOPC, 2% NiNTA (+iLID bound), 2% MCC-PE (+Ras bound), 2% $PIP_2$ lipids, +/- GβGγ. Unbound GβGγ was washed out of sample chamber prior to flowing in 2 nM AF647-PI3Kγ, 20 nM AF546-Btk, and where relevant, 5 nM PTEN and/or 50 nM GEF(no feedback). **(d)** Ras(GTP) and $PIP_3$ can spontaneously form bistable patterns in the presence of positive feedback and RasGAP inhibition. Representative TIRF-M images showing PI3Kγ-dependent $PIP_3$ generation colocalizing with regions of spontaneous Ras activation in the absence of 488 nm activating light. Membrane composition: 92% DOPC, 2% NiNTA (+iLID bound), 2% MCC-PE (+Ras bound), 4% $PIP_2$ lipids, GβGγ-farnesyl. Reaction contained: 30 nM GEF(fb), 20 nM GEF-SspB(micro), 690 nM RasGAP, 2 nM PTEN, 2 nM PI3Kγ, 20 nM AF546-LBD, and 50 nM AF647-RBD.

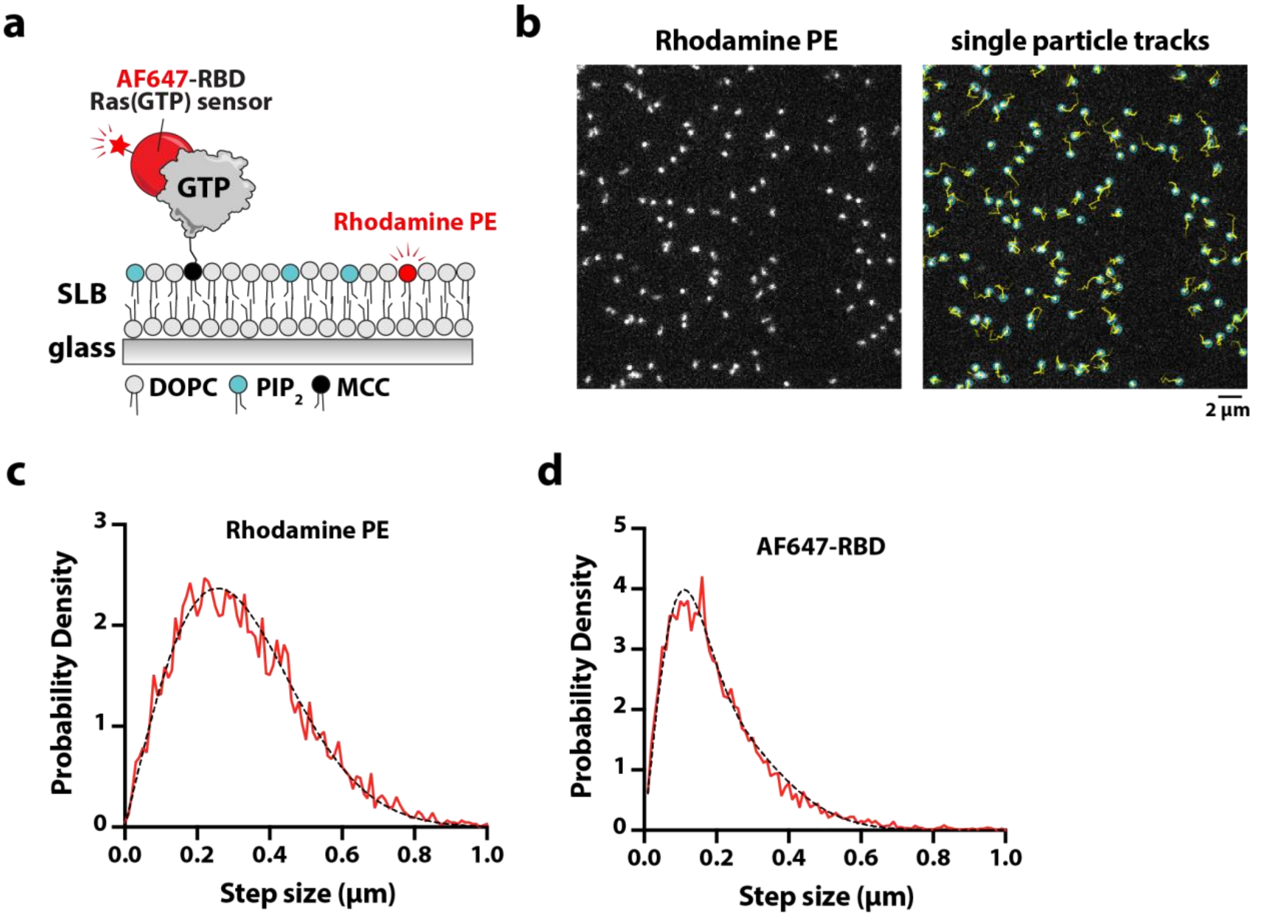


**Supplementary Figure 8**

**Ras(GTP) and lipid diffusion coefficients measured by single particle tracking**

**(a)** Experimental design for visualizing Ras(GTP) and lipid diffusion in supported lipid bilayers using single molecule TIRF microscopy. **(b)** Representative TIRF-M image showing the localization of lissamine rhodamine *phosphatidylethanolamine* (Rhodamine PE). Single particle detection (light blue circle) and trajectories (faded yellow lines) show dynamic lateral diffusion of lipids. **(c-d)** Step size distributions based on frame-to-frame single molecule displacements (µm) measured in the presence of **(c)** 10 pM AF647-RBD or (d) 0.00001% molar fraction of Rhodamine PE. Dashed black line represents the curve fit used to calculate the diffusion coefficient for **(c)** AF647-RBD (D1 = 0.27 µm$^2$/sec, D2 = 1.13 µm$^2$/sec, $\alpha$ = 0.57, n = 4922 events) and **(d)** Rhodamine PE (D = 2.76 µm$^2$/sec, n = 6361 events). For the two-component fit of the AF647-RBD step size distribution, alpha ($\alpha$) corresponds to the fraction of the single molecule events that are described by the slow diffusion state (i.e. D1). Membrane composition: 92% DOPC, 2% NiNTA (+iLID bound), 2% MCC-PE (+Ras bound), 4% $PIP_2$ lipids, GβGγ-farnesyl.

### SUPPLEMENTARY TABLE 1

| Parameter | 5d | 5d | 6h,i | 6h,i | 6h,i | 6h,i | S6 |
|---|---|---|---|---|---|---|---|
| L (grid width, pixels) | 100 | 100 | 100 | 100 | 100 | 100 | 100 |
| T (total # of time steps) | 500 | 500 | 2000 | 2000 | 2000 | 2000 | 1000 |
| r (growth rate) | 0 | 4 | 0.5 | 0.5 | 0.5 | 0.5 | 0 |
| D (diffusion, $\mu m^2/sec$) | 0.5 | 0.5 | 0.5 | 1 | 2 | 4 | 1, 3 |
| K (max Ras density per $\mu m^2$) | 20 | 20 | 100 | 100 | 100 | 100 | 100 |
| dx (step size is x dimension) | 1 | 1 | 1 | 1 | 1 | 1 | 1 |
| dy (step size is y dimension) | 1 | 1 | 1 | 1 | 1 | 1 | 1 |
| dt (time step, seconds) | 0.03 | 0.03 | 0.05 | 0.05 | 0.05 | 0.05 | 0.05 |
| decay rate (lambda, $\lambda$) | 3.1 | 3.1 | 0.25 | 0.25 | 0.25 | 0.25 | 0 |
| radius of perturbation (pixel) | 10 | 10 | 10 | 10 | 10 | 10 | 10 |
| Initial Ras density (per $\mu m^2$) | 0.5 | 0.5 | 25 | 25 | 25 | 25 | 25 |

Parameters for Fisher wave simulations. The corresponding figure numbers are printed at the top each column.
