## Supplementary File_FisherWave simulation for "Reconstitution of Ras-PI3Kγ membrane communication and feedback using light-induced signaling inputs"

```python
# -*- coding: utf-8 -*-
"""
This code is used to simulate a Ras(GTP) Fisher Wave on a 2-dimensional
membrane surface in the presence of global GAP-mediated inhibition

Manuscript title:
Reconstitution of Ras-PI3Kγ membrane communication and feedback using
light-induced signaling inputs

Authors:
Sophia Doerr (1,3), Andres Olavarrieta-Colasurdo (2,3,4), Scott D. Hansen (2,3)

(author affiliations):
1 Department of Biology, University of Oregon, Eugene, OR 97403
2 Department of Chemistry and Biochemistry, University of Oregon
3 Institute of Molecular Biology, University of Oregon, Eugene, OR 97403
4 Current address: University of California at San Francisco

"""

# %%
# Simulation of a Fisher Wave with a circular perturbation

import numpy as np
import matplotlib.pyplot as plt
import tifffile

#%%

# Parameters
L = 100               # Grid size (L x L) (pixels)
T = 1000              # Number of time steps (frames)
r = 4               # Intrinsic growth rate
D = 0.5             # Diffusion coefficient (µm2/sec)
K = 20              # Carrying capacity, Density of Ras molecules per µm²
dx = 1              # Grid spacing
dy = 1              # Grid spacing
dt = 0.03           # Time step
lambda_decay = 3.1  # Decay constant (rate at which u decays)

# Derived parameters
nx = int(L / dx)   # Number of grid points in x-direction
ny = int(L / dy)   # Number of grid points in y-direction
u = np.zeros((nx, ny))   # The population/concentration array
u_new = np.zeros((nx, ny))

# Radius of the circular perturbation
radius = 10  # Adjust the radius as needed

# Apply a circular perturbation at the center of the grid
for i in range(nx):
```

```
    for j in range(ny):
        # Calculate the distance from the center of the grid
        distance = np.sqrt((i - nx//2)**2 + (j - ny//2)**2)
        if distance <= radius:
            u[i, j] = 1
# Applies a circular perturbation that activates 5% of the
# maximum Ras(GTP) density

# Function to update the field u using the Fisher equation with decay
def update(u, r, D, K, dx, dy, dt, lambda_decay):
    u_new = u.copy()

    # Loop over the interior grid points
    for i in range(1, nx-1):
        for j in range(1, ny-1):
    # Laplacian approximation
    laplacian = (u[i+1, j] + u[i-1, j] + u[i, j+1] + u[i, j-1]
                 - 4 * u[i, j]) / (dx**2)

    # Fisher equation update with GAP-mediated decay term
    u_new[i, j] = u[i, j] + dt * ((D * laplacian) + (r * u[i, j]
                  * (1 - u[i, j] / K)) - (lambda_decay * u[i, j]))

    # Boundary conditions (set boundaries to zero)
    u_new[0, :] = u_new[-1, :] = 0
    u_new[:, 0] = u_new[:, -1] = 0

    return u_new

# Simulation loop

# Set the frequency of frames displayed and saved
frame_interval = 1  # Save every X time steps

for t in range(T):  # The loop iterates T times, where T is the total
# number of time steps (defined in parameters above)
    u = update(u, r, D, K, dx, dy, dt, lambda_decay)
    # Update the grid using the Fisher equation with decay
# This is the main simulation loop, where the variable t represents
# the current time step (ranging from 0 to T-1).
# For each iteration, the grid u is updated by calling the update()
# function, for the reaction diffusion system

    # Save every `frame_interval` time steps
    if t % frame_interval == 0:
        # This condition checks if the current time step t should be saved.
        # The modulus operator % is used to check the remainder when t is
        # divided by frame_interval.
        # If t % frame_interval == 0, it means that the current time step
        # is a multiple of frame_interval. then frame is saved.
        # Save directly to TIFF stack (append mode)
```

```
        if t == 0:
            tifffile.imsave('/write path here/output_FR0.tif', u)
            # First frame, create the file
        else: # For all subsequent time steps (t > 0),
              # this code saves subsequent frames
            tifffile.imsave(f"/write path here/output_ALL.tif", u, append=True)
            # All frames from simulation, create the file
# change the file path to save output in desired directory
# %%
```
